# Chromatin priming by transcription factor ELF3 confers transcriptional competence for amniotic differentiation on primed human embryonic stem cells

**DOI:** 10.1101/2025.08.04.668400

**Authors:** Masatoshi Ohgushi, Kaori Honda, Hitoshi Niwa, Mototsugu Eiraku

## Abstract

The self-organizing emergence of the amniotic lineage in synthetic human embryo models suggests that embryonic pluripotent cells possess an intrinsic potential to initiate extraembryonic amniotic differentiation. However, it remains poorly understood how this intrinsic amniotic competence is established and how it is linked to pluripotency. In this study, we found that the transcription factor ELF3, whose expression is restricted under primed pluripotency, governs the availability of amniotic competence. ELF3 is induced upon perturbation of primed pluripotency *in vitro* and expressed in cell populations associated with extraembryonic competence *in vivo*. We show that ectopic expression of ELF3 in primed pluripotent stem cells activates an amniotic transcriptional program, while genetic ablation of ELF3 impairs induction of amnion-related genes in an extraembryonic differentiation model. Mechanistically, ELF3 binds cis-regulatory elements of extraembryonic and amnion-related genes and increases their accessibility, thereby facilitating histone modifications, transcriptional cofactor recruitment, and transcriptional remodeling toward amniotic differentiation, including autocrine BMP-dependent signaling. Using a BMP4 enhancer as a model, we show that the ELF3-mediated increase in chromatin accessibility is required for TEAD/TAZ recruitment and subsequent transcriptional induction. These findings indicate that ELF3-mediated chromatin priming enables the retrieval of latent amniotic competence embedded within primed pluripotency, offering a new perspective on the developmental regulation of lineage plasticity in early human embryogenesis.

## Introduction

Embryogenesis exhibits substantial diversity across species in embryonic morphology, tissue allocation, molecular signatures, and the mechanisms and timing of fate specification. Over the past few decades, mice have served as the predominant model for studying mammalian development, providing valuable insights into human embryogenesis. However, notable differences between mouse and human embryogenesis have come to light (*1*). Among them, the development of the amniotic ectoderm, a squamous epithelial tissue that gives rise to the future amnion, has emerged as a particularly intriguing process (*1, 2*). Although this lineage originates from pluripotent epiblasts in both species, its formation processes differ markedly, folding in mice versus cavitation in humans (*3*). Moreover, this lineage segregates shortly after implantation in humans, whereas in mice it emerges only after gastrulation onset (*2, 4, 5*). In addition to these morphogenetic and temporal differences, recent studies have highlighted primate-unique functions of the amniotic ectoderm in primordial germ cell specification and gastrulation initiation (*6,7*). Thus, the amniotic ectoderm serves not only as the origin of the amnion but also as a functionally critical extraembryonic tissue in human development. Despite such significance, many aspects of its development and function remain incompletely understood, in part due to the limited accessibility of peri-implantation stage human embryos (*8–12*). As an alternative approach, human pluripotent stem cells (PSCs) have been widely used to model amniogenesis *in vitro* (*13–26*), and this has, in turn, brought to light emerging questions.

First, amniotic ectoderm-like cells can be induced not only from naïve PSCs but also from primed PSCs, raising an open question of why primed PSCs, aligned with a post-implantation epiblast stage that follows amniotic emergence (*19, 27, 28*), can still undergo amniotic differentiation. This temporal mismatch suggests that primed PSCs may retain an unexpected degree of epigenetic plasticity, although this remains hypothetical. Second, the regulatory logic initiating lineage segregation from the pluripotent epiblast has come into question. The canonical views, primarily established through genetic studies in mice, posits that extraembryonic tissues, particularly trophoblast cells, provide instructive cues for amniotic differentiation primarily via the secretion of BMP ligands (*29, 30*). Mimicking this paradigm, many PSC-based systems have employed BMP ligands to induce amniotic differentiation (*15, 16, 19–21, 24–26*). However, recent progress in generating synthetic human embryos provided the possibility that amniotic ectoderm-like structures arise in the absence of exogenous BMP or BMP-producing trophoblast lineages (*31–33*). In combination with other studies showing the cell-autonomous emergence of amnion-like cells was also observed in other stem cell models (*13, 14, 17, 18, 20, 22*). Notably, such a cell-intrinsic mode of amnion specification may also resonate with the evolutionary view that the amnion predates the placenta and therefore may have originated through a trophoblast-independent developmental program (*3, 34*). Although these observations suggest that human PSCs possess a latent competence to initiate amniotic differentiation independent of external instructive inputs, how such an intrinsic differentiation competence becomes available in primed pluripotent cells remains unresolved.

In this study, we identify ELF3 as a transcription factor whose ectopic expression remodels global transcriptome of human primed embryonic stem cells (ESCs) to an amnion-like state. We demonstrated that chemical perturbation of primed pluripotency leads to ELF3 upregulation, which is required for the subsequent induction of a series of amniotic-related genes. Mechanistic analyses using BMP4 regulation as a model show that ELF3 facilitates chromatin accessibility at cis-regulatory elements of amniotic genes, thereby priming these elements for the recruitment of transcriptional regulators and general transcriptional cofactors that promote transcription. These findings suggest a chromatin-priming role of ELF3 that is constrained under primed pluripotency, thereby providing insight into a broader principle by which latent differentiation potentials embedded in primed pluripotent states can be reactivated through intrinsic programs.

## Results

### Screening the putative regulator of embryonic-to-extraembryonic transition

We previously reported that using small-molecule inhibitors A83-01 (targeting ACTIVIN/NODAL signaling) and PD173074 (targeting FGF signaling), referred to hereafter as an ‘AP’, human ESCs were efficiently converted into GATA3^+^ extraembryonic cells (ExECs) (*22*). The AP treatment led to the downregulation of pluripotency-associated genes and the induction of multiple genes associated with extraembryonic phenotypes (Fig. 1A and Fig. S1A-B). Our previous single-cell RNA sequencing (scRNA-seq) study indicated that AP-treated cells represent a heterogeneous extraembryonic population, including amnion-like and syncytiotrophoblast (STB)-like cells (Fig. S1C) (*22*). To understand the molecular basis of extraembryonic competence in primed ESCs, we first sought to identify key transcription factors (TFs) that drive the ESC-to-ExEC transition. To monitor the transition to extraembryonic state in a non-invasive manner, we utilized a GATA3 reporter ESC line, G3KI#2-5, in which a tdTomato (tdT) fluorescence protein is knocked into the *GATA3* locus (*22*). To profile gene expression dynamics during the first 2 days of AP-induced extraembryonic conversion, we performed bulk mRNA-sequencing (mRNA-seq) analyses. This identified 1,874 differentially expressed genes (DEGs), including 898 upregulated and 976 downregulated genes (false discovery rate (FDR) < 0.05, p < 0.01, [log_2_ fold-change] > 2 or < -2) (Fig. 1B). Among the upregulated DEGs, 53 encoded TFs, including canonical BMP target genes such as *ID1, ID2, ID4*, *SMAD6*, *MSX1,* and *MSX2*, suggesting activation of endogenous BMP signaling, as we previously demonstrated (*22*). To distinguish primary transcriptional responses to AP from secondary effects mediated by endogenous BMP signaling, we compared the expression profiles of the 53 TFs in cells treated with AP alone or with AP plus BMP blockers (LDN-193189 and recombinant Chordin). Hierarchical clustering analysis separated these TFs into BMP-dependent and BMP-independent groups (Fig. 1C). Strikingly, the majority of AP-induced TFs (42 out of 53), including key regulators and markers associated with extraembryonic states, *GATA3, TFAP2A, CDX2, ISL1, TBX3, MSX2, SP6,* and *HAND1*, were secondarily induced via endogenous BMP signaling (Fig. 1C).

**Fig. 1.**
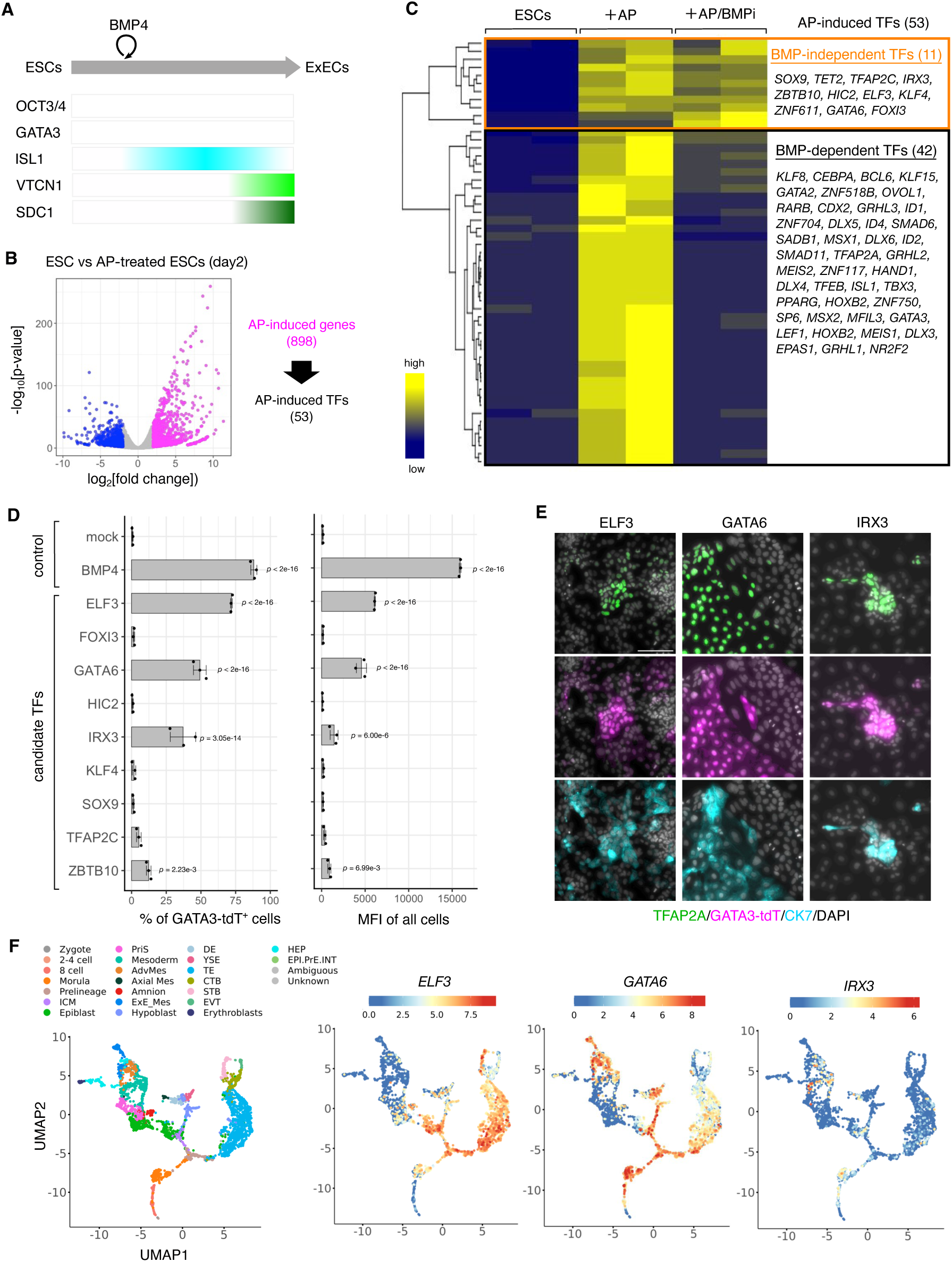
Search for transcription factors capable of inducing GATA3^+^ ExECs. (**A**) Schematic diagram of marker gene expression kinetics during AP-induced differentiation. (**B**) Volcano plot comparing GATA3^+^ ExECs (day 2) with ESCs. Two biological replicates per condition. Upregulated and downregulated genes are indicated as orange and blue dots, respectively. From the 898 upregulated genes, 53 genes encoding transcription factors were selected for subsequent analyses. (**C**) Heatmap showing expression of 53 TFs in ESCs (left), AP-treated cells (2 days, center), and AP-treated cells in the presence of BMP blockers (2 days, right). Two biological replicates per condition. Gene expression is shown as scaled TPMs for each replicate. **(D**) Percentage of GATA3-tdTomato^+^ cells (left) and mean fluorescence intensity (MFI) across all cells (right). ESCs engineered to express the indicated transgenes were treated with dox for 4 days. Data are presented as mean ± SD (n = 3 independent experiments). Statistical significance was determined by Dunnett’s test versus mock. (**E**) Immunostaining of extraembryonic markers in the cells expressing *ELF3* (left), *GATA6* (center), and *IRX3* (right). Scale bar, 100 µm. Two independent cultures were used for the experiment. **(F**) UMAP projection of human embryo scRNA-seq datasets and expression profiles of *ELF3*, *GATA6*, and *IRX3* mRNA on the UMAP. Each dot is colored by the cell annotations as defined in the original publication.

Based on the hypothesis that upstream activators of BMP signaling would be included in the BMP-independent group, we selected nine TFs from this group for further analyses. We cloned the corresponding genes and tested whether their ectopic expression could induce tdT fluorescence in G3KI#2-5 ESCs. To achieve controlled expression, we employed a doxycycline (dox)-inducible transgene system and established polyclonal ESC populations capable of expressing each candidate TF upon dox supplementation (Fig. S2A). In parallel, we generated BMP4-inducible ESCs as a positive control and G418-resistant gene-expressing ESCs as a negative control. As expected, BMP4-inducible cells, but not negative control cells, showed tdT expression upon dox supplementation, validating the functionality of our drug-controllable expression system (Fig. 1D and Fig. S2B). Through this screening, we identified three TFs, ELF3, GATA6, and IRX3, that significantly increased tdT fluorescence (p < 0.001) (Fig. 1D). Among these, ELF3 induced markedly higher tdT expression, whereas IRX3 elicited moderate increases in only a subset of cells (Fig. S2B). GATA6-expressing cells displayed a bimodal distribution of tdT fluorescence, with populations showing high and moderate levels (Fig. S2B). Immunostaining analyses further confirmed that tdT-positive cells induced by these TFs co-expressed TFAP2A and CK7, both recognized as markers of extraembryonic differentiation (Fig. 1E).

To assess the relevance of these three candidate TFs to extraembryonic phenotypes, we analyzed their expression dynamics throughout human embryogenesis, from the zygote stage to gastrulation, using a publicly available integrated human embryonic scRNA-seq dataset (*35*). This analysis revealed distinct mRNA expression profiles of three candidates (Fig. 1F and Fig. S2C). *GATA6* is broadly expressed across multiple extraembryonic lineages but is low in amniotic populations. *IRX3* expression is less prominent in all extraembryonic cells. Interestingly, *ELF3* is broadly abundant in populations that possess extraembryonic competence and in trophoblastic and amniotic lineages, implying its possible roles in maintaining extraembryonic competence or identity.

Based on these observations, we regarded ELF3 as a promising candidate associated with the embryonic-to-extraembryonic transition and conducted further functional analysis.

### Characterization of the transcriptional changes upon ELF3 induction

To comprehensively characterize cellular responses to ELF3 induction, we established two independent HA-ELF3-inducible ESC clones from the drug-resistant pool (tet-ELF3#3 and tet-ELF3#5) (Fig. 2A and Fig. S3A). Although these clones exhibited different levels of ELF3 induction upon dox supplementation, they showed consistent GATA3-tdT fluorescence levels and induction kinetics (Fig. S3A-B). Hereafter, we primarily present data obtained from clone tet-ELF3#3. We confirmed that ELF3 expression promotes transcription of *BMP4* and BMP-dependent TFs (Fig. S3C-F), supporting our hypothesis that ELF3 functions upstream of BMP signal activation. Live imaging and flow cytometric analyses further indicated that the majority of dox-treated cells (> 95%) co-expressed GATA3 and extraembryonic surface marker APA (Fig. 2B-C). Immunostaining demonstrated a gradual attenuation of pluripotency-associated proteins, concomitant with the upregulation of several genes associated with extraembryonic phenotypes (Fig. S3G). These results indicate that ELF3 expression alone can drive two distinct outcomes: the disruption of pluripotency and the induction of extraembryonic features.

**Fig. 2.**
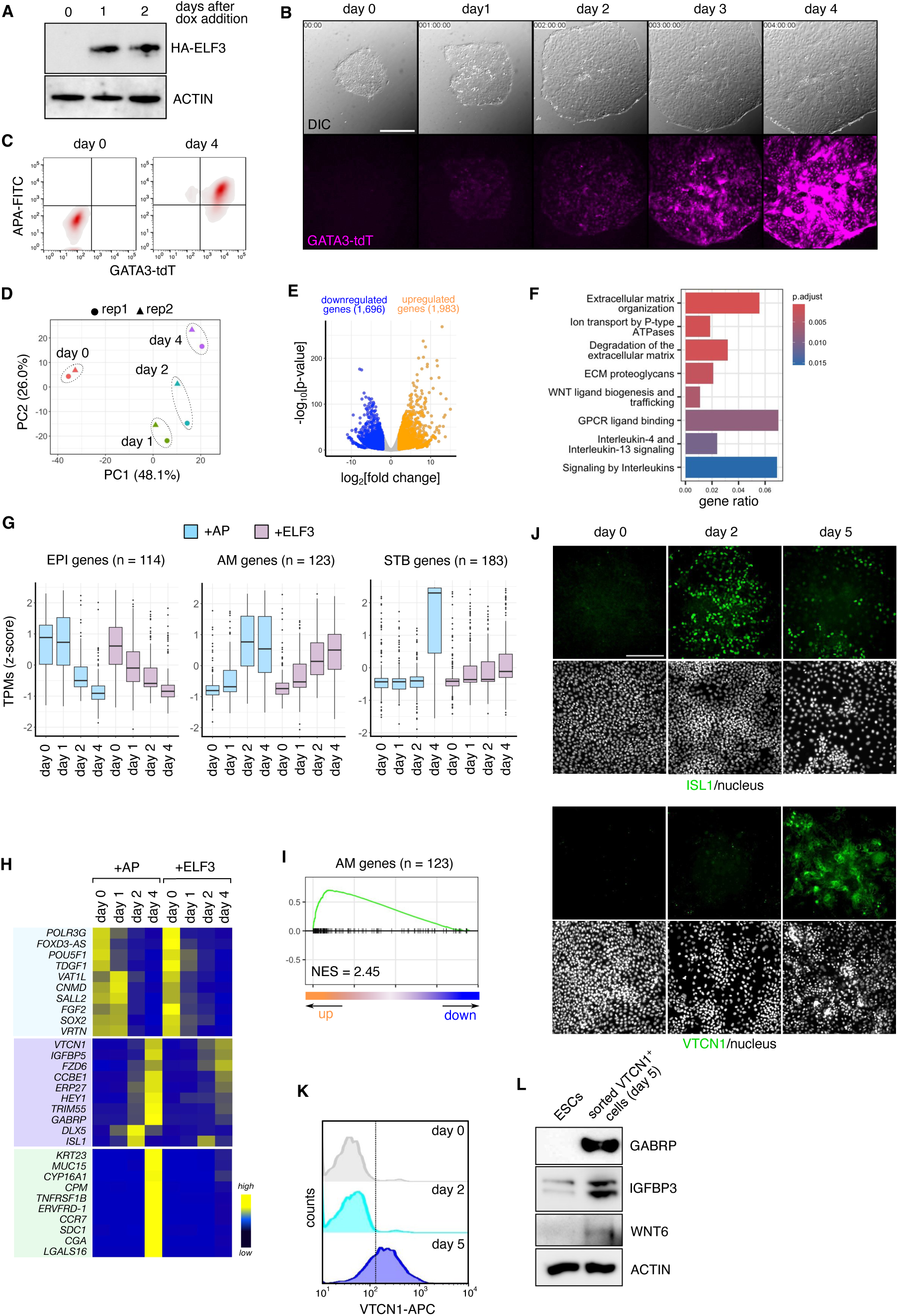
Ectopic expression of ELF3 promotes transition to an amnion-like state. (**A**) Representative western blot (WB) images showing dox-mediated induction of HA-tagged ELF3 from two independent experiments. ACTIN as a loading control. (**B**) Live imaging of tet-ELF3#3 ESCs treated with dox for 72 h. Bottom panels show GATA3-tdTomato fluorescence (magenta). Recording started immediately after dox addition. Three experiments were done. Scale bar, 500 µm. (**C**) Flow cytometry of APA^+^/GATA3^+^ cells at day 0 or day 4 after dox addition. Representatives are shown from three independent experiments. (**D**) PCA of ELF3-induced transcriptome changes (two biological replicates per condition). The top 1,000 variable genes were used for calculation. (**E**) A volcano plot comparing ELF3-expressing cells (dox, 4 days) versus control ESCs. Upregulated and downregulated genes are indicated as orange and blue dots, respectively. (**F**) Gene ontology analyses of upregulated genes. (**G**) Boxplot showing expression dynamics of epiblast genes (EPI; n = 114), amnion (AM; n = 123), and STB (n = 183) genes in the cells treated with dox for the indicated days. Marker genes defined by published human embryo scRNA-seq data (Xiang et al., 2020, Tyser et al, 2021, Ohgushi et al, 2022). Data are means of scaled TPMs from two replicates. The center lines in each box indicate the median. AP-treated cultures are included as a control reference. (**H**) Heatmap showing representative epiblast, amnion and STB marker genes in AP-treated cells and dox-treated tet-ELF3#3 cells (day 0, 1, 2, and 4). Amnion markers are highlighted (light pink box), epiblast markers (violet box), and STB markers (green box). Data are means of scaled TPMs from two replicates. (**I**) GSEA of ELF3-expressing cells (dox, 4 days) versus control ESCs using the same geneset as in **H** (NES; normalized enrichment score). (**J**) Immunostaining of the indicated amnion markers at the indicated time points after dox supplementation. Three independent cultures were used for the experiment. Scale bar, 100 µm. (**K**) Flow cytometry of VTCN1-expressing cells at day 0, 2, and 5 after dox addition. Representatives are shown from three independent experiments. (**L**) Representative WB images confirming the expression of amnion markers in the sorted VTCN1^+^ cells. AP-induced differentiation and flow cytometry sorting were done in two independent experiments. Lysates from ESCs were examined as an undifferentiated control. Actin was examined as a loading control.

To further dissect the genome-wide transcriptional changes induced by ELF3, we performed bulk mRNA-seq analyses. Principal component analysis revealed that global transcription was drastically remodeled within 24 hours after ELF3 induction (Fig. 2D). Notably, the immediately upregulated genes included *SPRR2A* and *CLDN4*, both previously reported as ELF3 target genes (Fig. S3H) (*36, 37*). At day 4, 1,983 were significantly upregulated and 1,696 genes downregulated (FDR < 0.05, p < 0.01, [log_2_ fold-change] > 2 or < -2) (Fig. 2E). To gain functional insight into the upregulated genes, we performed pathway enrichment analysis and found enrichment in biological pathways involved in extracellular matrix (ECM) remodeling, ion transport, and interleukin signaling, all implicated in the functions and activities of the extraembryonic membrane system (Fig. 2F).

Considering our previous results that primed ESCs could give rise to both amnion-like and STB-like cells (*22*), we then assessed expression patterns of a panel of lineage-associated genes, which were defined as highly expressed genes in the epiblast, amniotic ectoderm, or STB lineages of implantation-and gastrulation-stage human embryos (Fig. S3I-J) (*8, 11, 22*). These analyses revealed a clear increase in the amnion signature upon ELF3 induction, accompanied by a concurrent decline in epiblast identity (Fig. 2G). Remarkably, *ISL1* was upregulated as early as day 2 following ELF3 induction, whereas the induction of *VTCN1* and *GABRP* required an extended culture period (Fig. 2H). Together with the expression analyses, gene set enrichment analyses (GSEA) provided transcriptome-wide evidence for coordinated enrichment of the amniotic gene program following ELF3 induction (Fig. 2I). Consistent with these transcriptomic analyses, we also confirmed induction of amnion-associated proteins (Fig. 2J and Fig. S3K). Notably, a substantial proportion of dox-treated cultures were positive for VTCN1, and the VTCN1-positive fraction also contained additional amnion-associated proteins (Fig. 2K-L). Among these, VTCN1 and GABRP proteins have been reported in the amniotic ectoderm of developing primate embryos (*7, 15, 38*). On the other hand, most of the canonical STB-related genes, such as *SDC1*, *CGA*, and *ERVFRD-1*, remained low in the ELF3-induction setting (Fig. 2G-H), and consistently, we did not observe a robust induction of SDC1^+^ cells (Fig. S3L). These results revealed that ectopic expression of ELF3 promotes transcriptome remodeling suggestive of amnion-dominant extraembryonic differentiation.

### Loss-of-function validation of ELF3 in amniotic gene regulation

Given that ectopic ELF3 expression per se triggered gene expression changes reminiscent of amniotic differentiation, we next used a loss-of-function approach to corroborate the requirement of ELF3 in amniotic gene activation. To this end, we utilized the AP-induced differentiation model, in which a VTCN1^+^/GABRP^+^ amnion-like population arises under defined conditions (Fig. S1C). *ELF3* expression was immediately initiated upon AP-treatment and sustained throughout the subsequent culture period (Fig. 3A-B and Fig. S5A). Immunostaining analyses revealed that ELF3 protein was induced during ExEC differentiation (Fig. 3C). To evaluate the functional requirement of ELF3, we generated ELF3-knockout ESCs using the G3KI#2-5 cell line as a parental background (Fig. S5B). Two independent clones were established by excising the genomic region spanning exons 2 to 7 of the *ELF3* gene (E3KO#21 and #24) (Fig. S5C). Western blotting confirmed that ELF3 protein was undetectable in both E3KO clones (Fig. 3D). Upon AP treatment, these clones showed GATA3-tdT expression levels comparable to those of parental cells (Fig. S5D), suggesting that they retained extraembryonic differentiation potential. We then focused on the effect of ELF3 deficiency on AP-induced upregulation of VTCN1 and GABRP, representative genes whose transcripts are abundant in amnion-containing population in the human embryo scRNA-seq datasets (Fig. S5E). Quantitative PCR (qPCR) analyses revealed that ELF3 deficiency significantly suppressed their transcriptional induction (Fig. 3E). Consistently, immunostaining analyses confirmed lower levels of VTCN1 in AP-treated E3KO clones at the protein level (Fig. 3F). Inducible HA-ELF3 add-back during AP treatment increased VTCN1 protein levels in the knockout background compared with AP alone (Fig. S5F-H), indicating that the reduced expression of amnion-related genes in ELF3 KO cells is attributable to loss of ELF3.

**Fig. 3.**
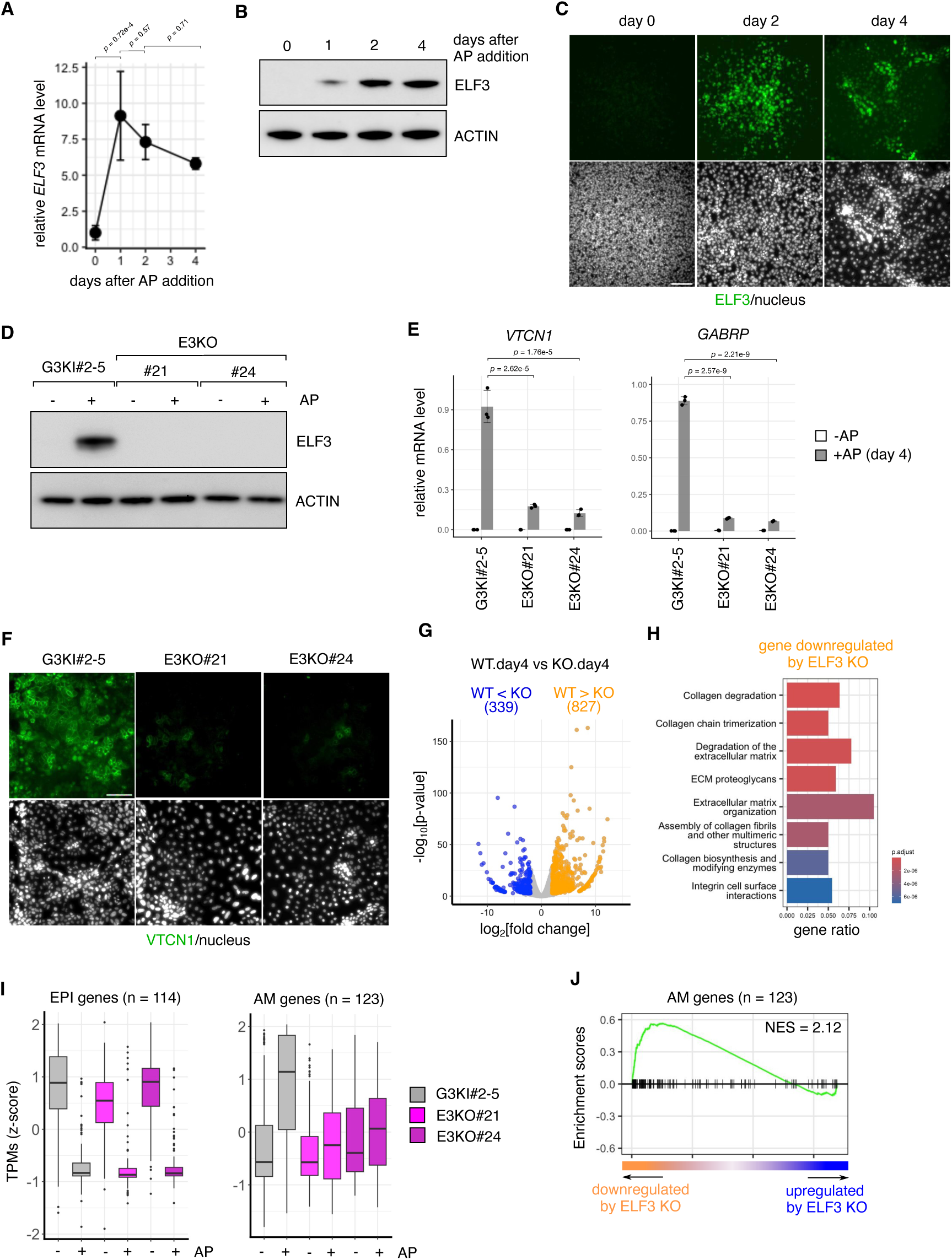
ELF3 deficiency impaired amniotic gene activation. (**A**) qPCR analysis of *ELF3* expression during AP-induced differentiation. Data are shown as relative to day 0, with mean ± SD (n = 3 independent experiments). Statistical significance was determined by Tukey’s test. (**B**) Representative WB images showing ELF3 expression dynamics during AP-induced differentiation from three independent experiments. Actin was used as a loading control. (**C**) Immunostaining of ELF3 in differentiating cells. Three independent experiments were done. Scale bar, 100 µm. (**D**) Representative WB images showing ELF3 expression in ELF3 knockout clones (#21 and #24) and parental G3KI#2-5 ESCs (two independent experiments). Cells were treated or left untreated with AP for 2 days. Actin was used as a loading control. (**E**) qPCR analysis of *VTCN1* and *GABRP* expression in ELF3-knockout clones and parental ESCs. Cells were treated or left untreated with AP for 4 days. Data are shown as arbitrary units with mean ± SD (n = 3 independent experiments). Statistical significance was determined by Dunnett’s test (vs AP-treated parental cells). (**F**) Immunostaining of VTCN1 in the indicated cells after AP treatment for 5 days. Three independent experiments were done. Scale bar, 100 µm. (**G**) Volcano plot comparing AP-treated ELF3-knockout cells with AP-treated parental ESCs (4 days). Genes downregulated (WT > KO) and upregulated (WT < KO) by ELF3 knockout are shown in orange and blue, respectively. (**H**) Gene ontology analyses of genes downregulated in ELF3-knockout cells. (**I**) Boxplot showing expression levels of amnion- and epiblast-related genes (as defined in Fig. 2g) in AP-treated (4 days) or untreated cells. Data are shown as scaled TPMs from two independent knockout clones and one parental cell line. The center lines in each box indicate the median. (**J**) GSEA of AP-treated ELF3-knockout versus AP-treated parental ESCs (4 days), using the same amnion and epiblast gene sets as in Fig. 2I.

To extend our analysis beyond a few representative markers, we performed bulk mRNA-seq analyses. The parental-vs-E3KO comparison of AP-treated cells at day 4 identified 827 genes whose upregulation was significantly diminished upon ELF3 deficiency (FDR < 0.05, p < 0.01, [log_2_ fold-change] > 1 or < -1), including several amnion-related genes (Fig. 3G and Fig. S5I). Pathway enrichment analyses of these downregulated genes revealed profound defects in ECM remodeling activity in ELF3-knockout cells (Fig. 3H). We further characterized the transcriptome changes in AP responses upon ELF3 deficiency using lineage-relevant gene sets defined from *in vivo* epiblast and amnion. This analysis demonstrated that although AP treatment caused a reduction in the epiblast signature in both parental and E3KO clones, the upregulation of the amniotic signature was substantially diminished in E3KO cells (Fig. 3I). Consistently, GSEA supported the attenuation of the amnion-associated transcriptional program in AP-treated E3KO cells (Fig. 3J). These results demonstrated that ELF3 is required for robust induction of an amnion-associated transcriptional program under AP conditions.

### Genome-wide profiling of ELF3 binding

To understand how ELF3 remodels the transcriptional landscape, we utilized the dox-inducible system and performed anti-HA chromatin immunoprecipitation followed by sequencing (ChIP-seq) to profile genome-wide ELF3 binding sites. As a 2-day treatment was sufficient to induce irreversible extraembryonic conversion of ESCs (Fig. S6A), we chose this time point for ChIP analyses. ELF3 ChIP-seq identified 31,103 genomic loci significantly bound by ELF3 (*q* < 1 x 10^-10^; compared with input), predominantly mapped to promoters, introns, or intergenic regions (Fig. 4A-C and Fig. S6B). Among the ELF3-regulated genes, 17.4 % (286 out of 1,644) of upregulated and 2.7 % (32 out of 1,201) of downregulated DEGs harbored ELF3 binding peaks within 2 kilobases of their transcription start sites (TSSs) (Fig. S6C). To investigate relevant histone marks in proximity to the identified ELF3-bound loci, we conducted ChIP-seq analyses for representative histone modifications, including histone H3 lysine 4 trimethylation (H3K4me3), lysine 27 trimethylation (H3K27me3), and lysine 27 acetylation (H3K27ac) (Fig. 4D-E). ELF3-bound genomic regions were typically devoid of H3K27me3, with no substantial change in H3K4me3 deposition, while accumulation of H3K27ac, a histone mark associated with active promoters and enhancers, was evident at a substantial proportion of ELF3 binding sites. In line with these observations, upregulated DEGs showed prominent enrichment of both ELF3 and H3K27ac peaks (Fig. 4F). Indeed, ELF3 and H3K27ac peaks frequently overlapped at putative ELF3 target gene loci (Fig. 4G and Fig. S6D). When focusing on gene loci associated with amnion phenotypes, we confirmed that ELF3 binding was detected at a substantial fraction of these gene loci, accompanied by an increase in H3K27ac signals (Fig. S6E). In contrast, several pluripotency-related genes downregulated upon ELF3 induction showed a remarkable reduction in active histone marks, despite the absence of clear ELF3 binding (Fig. S6F). These results suggested that ELF3 binds to cis-regulatory elements of amnion-related genes and is associated with their transcriptional activation, whereas downregulation of pluripotency-related genes occurs indirectly as a secondary consequence of ELF3 induction.

**Fig. 4.**
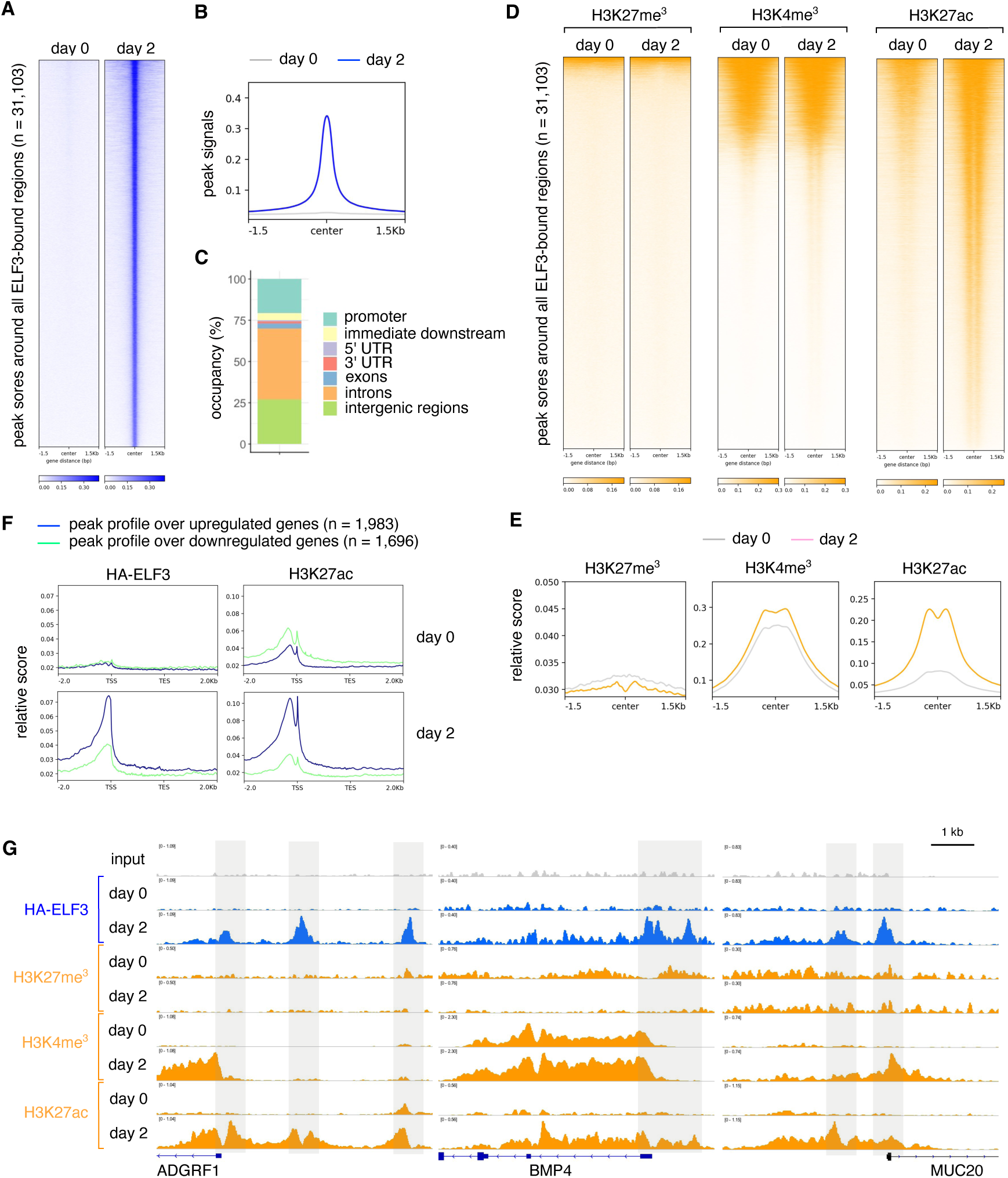
Profiling of ELF3 binding and histone modifications. (**A-B**) Coverage heatmaps (A) and profiles (B) showing HA-ELF3 binding peaks in dox-treated (2 days) and untreated cells. (**C**) Genomic distribution of HA-ELF3 binding peaks. (**D-E**) Coverage heatmaps (D) and profiles (E) showing H3K27me3 (left), H3K4me3 (center), and H3K27ac (right) in dox-treated (2 days) and untreated cells around ELF3-bound genomic loci (as defined in A). (**F**) Coverage profiles of HA-ELF3 (left) and H3K27ac (right) around genes upregulated (blue lines) or downregulated (green lines) in dox-treated (2 days) and untreated cells. (**G**) Genomic tracks showing the normalized ChIP-seq signals at the indicated genes.

### Chromatin accessibility remodeling upon ELF3 induction

The binding of TFs to cis-regulatory elements is often coupled with dynamic changes in chromatin accessibility. To investigate whether ELF3 influences chromatin accessibility, we performed the assay for transposase-accessible chromatin using sequencing (ATAC-seq) to profile genome-wide chromatin accessibility changes following ELF3 induction (*39*). We first examined the accessibility of ELF3-bound sites before induction and found that most ELF3 binding occurred at regions with low chromatin accessibility in the undifferentiated state (90.8 % of all ELF3 binding, Fig. S7A). We then identified differentially accessible regions (DARs) that either gained or lost accessibility following ectopic ELF3 induction (Fig. 5A). Notably, only a small subset of regions exhibited decreased accessibility, indicating that ELF3 primarily promotes chromatin opening (Fig. S7B). 7,689 regions, representing approximately 37 % of all open DARs, overlapped with ELF3 binding peaks (Fig. 5B and Fig. S7C-D). To explore whether an ELF3-dependent increase in chromatin accessibility is associated with recruitment of general transcriptional cofactors, we examined ELF3-bound accessible regions for occupancy by the histone acetyltransferase p300 and the Mediator complex, two cofactors previously implicated in ELF3-mediated transcription (*40, 41*). Both p300 and Mediator were enriched at these regions following dox treatment (Fig. 5C-D). At genomic regions proximal to ELF3-induced genes such as *SPRR2A*, *VGLL1*, *GATA3*, and *PRTG,* we observed overlapping ELF3 binding, p300 and Mediator recruitment, increased H3K27ac deposition, and elevated chromatin accessibility (Fig. 5E and Fig. S7E). For instance, at *VGLL1* loci, ELF3 binding was associated with enhanced accessibility immediately upstream of the TSS, annotated as a promoter (Fig. 5E). Additionally, ELF3 binding and chromatin opening were observed approximately 1.5 kilobases upstream of the *SPRR2A* TSS (Fig. 5E), suggesting that this region may function as an enhancer. Chemical inhibition of p300 acetyltransferase activity reduced the induction of these genes, confirming the role of p300 in their transcriptional regulation (Fig. S7F-G). Importantly, motif enrichment analysis of all ELF3-bound, accessibility-gained DARs revealed an overrepresentation of binding motifs for TFs known to be active in amnion-associated contexts, including GATA, TEAD, and TFAP2 factors (Fig. 5F). These observations suggest that ELF3 remodels a cis-regulatory accessibility landscape to initiate amnion-associated gene programs.

**Fig. 5.**
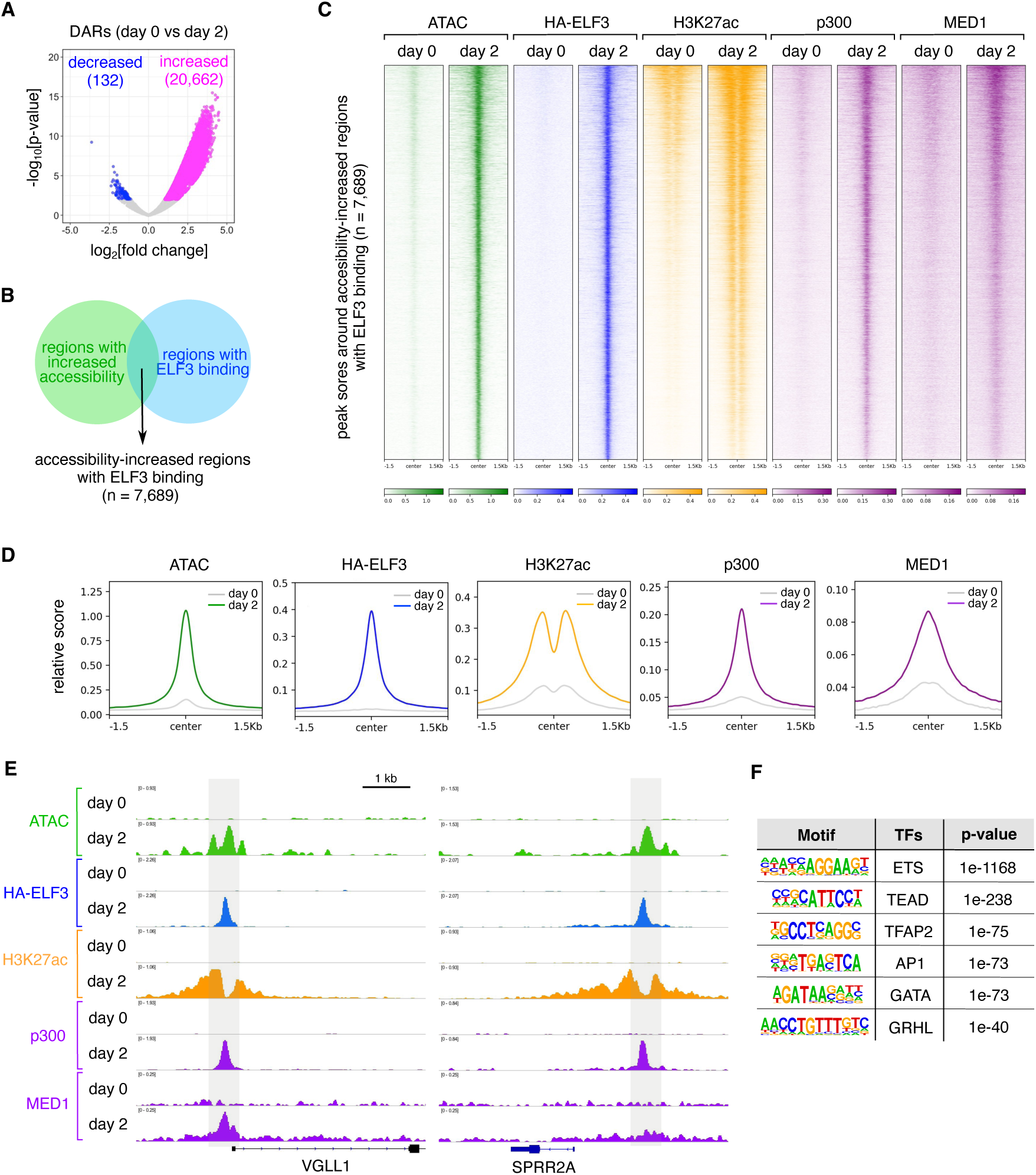
Chromatin accessibility dynamics of ELF3-binding genomic regions. (**A**) Volcano plot of differentially accessible regions comparing dox-treated (2 days) and control cells. Two replicates of ATAC-seq data were used for statistical calculation. Regions with increased or decreased accessibility are shown as orange and blue, respectively. (**B**) Venn diagram showing overlap between regions with increased accessibility and HA-ELF3-bound regions. (**C-D**) Coverage heatmaps (C) and profiles (D) of ATAC, HA-ELF3, H3K27ac, p300, and MED1 signals in dox-treated (2 days) and control cells around ELF3-bound regions with increased accessibility (as identified in B). (**E**) Genomic tracks showing the normalized ATAC-seq and ChIP-seq signals at the indicated loci. VGLL1 and SPRR2A are shown as representative ELF3-induced genes. (**F**) Transcription factor binding motifs enriched in ELF3-bound regions with increased accessibility. The top-ranked motif is an ETS site, consistent with ELF3 binding at these regions.

### Identification of BMP4 distal enhancers regulated by ELF3

Because BMP4 transcriptional activation represents one upstream event in ELF3-induced activation of amnion-associated genes (Fig. S3C-F), we examined the BMP4 locus as a model for ELF3-mediated control of cis-regulatory elements. The *BMP4* gene resides within a “gene desert”, lacking protein-coding genes within several hundred kilobases upstream or downstream. ELF3 binding and increased chromatin accessibility were observed at the *BMP4* TSS (Fig. 6A). In addition, we identified several ELF3-bound regions with enhanced accessibility located within 100 kilobases upstream and downstream of the *BMP4* TSS. We designated these regions BE1, BE2, BE3, and BE4 (Fig. 6A). BE1 was located near the TSS, whereas BE2-4 were found in distal regions downstream of the gene body. Among these ELF3-bound regions, BE2, BE3, and BE4, but not BE1, also displayed recruitment of p300 and accumulation of H3K27ac, supporting their potential role as active enhancers (Fig. 6A). To validate this hypothesis, we cloned each genomic region into luciferase reporter constructs and tested their transcriptional activity in ESCs (Fig. 6B). The BE1 element alone did not affect luciferase transcription (Fig. 6C). However, robust luciferase activation was observed when all four elements were linked in tandem (Fig. 6C), suggesting that at least one of BE2, BE3 or BE4 contributes to enhancer activity. To determine which elements were functionally essential, we removed each region from the reporter. Strikingly, deletion of BE3 caused a marked reduction in luciferase transcription (Fig. 6D). We further confirmed that a construct containing only BE3 was sufficient to drive robust luciferase activity (Fig. 6E). These results indicate that BE3 likely served as a major transcriptional enhancer in this context.

**Fig. 6.**
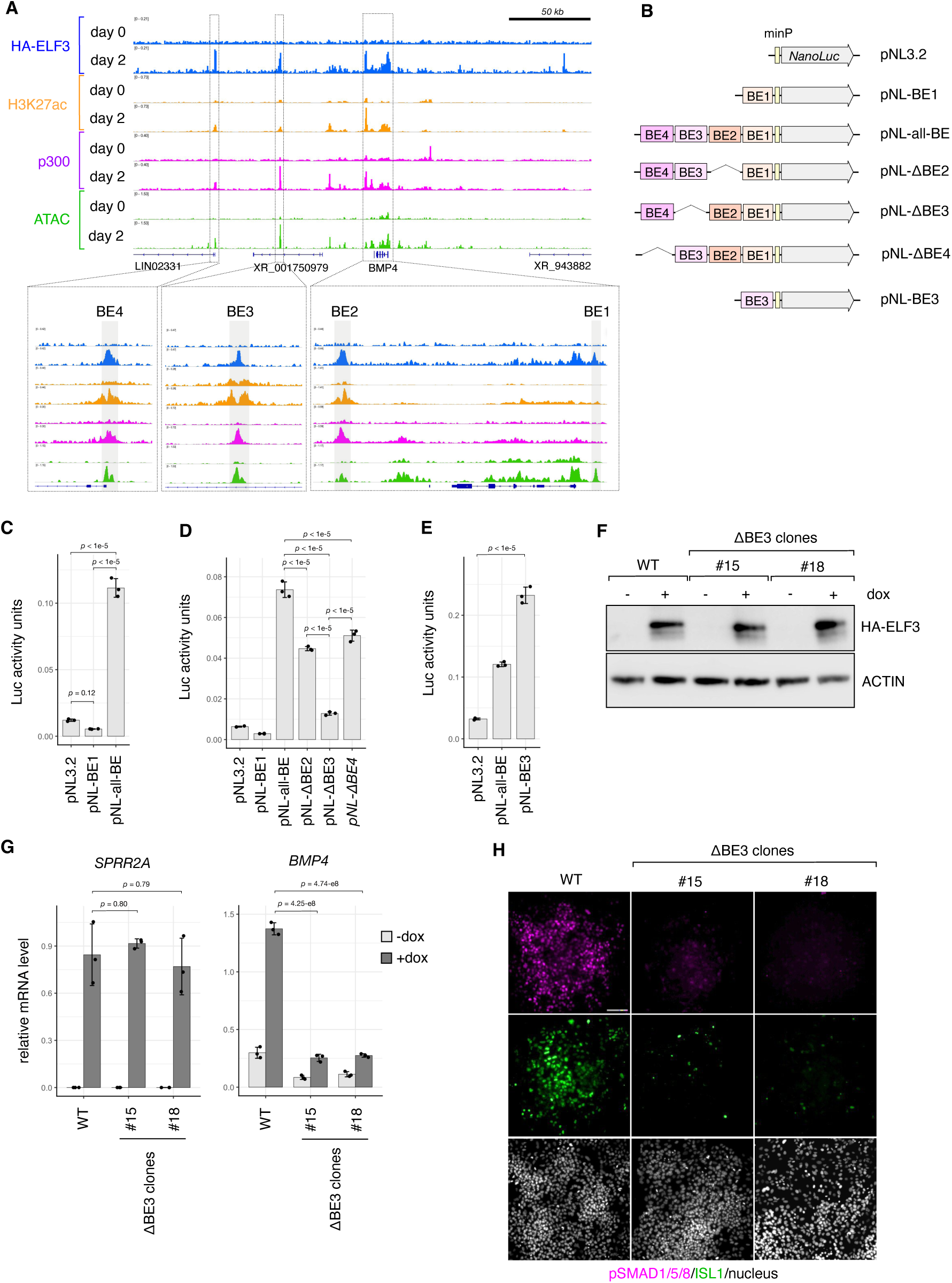
Characterization of ELF3-bound BMP4 distal enhancers. (**A**) Genome tracks showing co-enrichment of ELF3-binding, ATAC peaks and epigenetic hallmarks (H3K27ac, p300) around the BMP4 locus. Lower panels show enlarged views of the boxed regions in the upper image. ELF3 binding overlaps with ATAC peaks upstream (BE1) and downstream (BE2-4) of the BMP4 gene body. (**B**) Schematic of luciferase reporter constructs used in the assay. (**C-E)** Luciferase reporter assay showing enhancer activity of BE elements. Cells were transiently transfected with the indicated NanoLuc reporter constructs and an internal control reporter. NanoLuc activities was measured 48 h after transfection, normalized to the control, and plotted as mean ± SD (n = 3 independent experiments). Statistical significance was determined by Tukey’s test. (**F)** Representative WB images showing HA-ELF3 induction in ΔBE3 clones (#15 and #18) and parental ESCs (WT) from two independent experiments. Cells were treated or untreated with AP for 2 days. Actin was examined as a loading control. (**G**) qPCR quantification of *SPRR2A* and *BMP4* expression in ΔBE3 clones and parental ESCs (WT). Cells were treated or untreated with dox for 2 days. Data are plotted as mean ± SD (n = 3 independent experiments). Statistical significance was determined by Dunnett’s test (vs dox-treated parental cells). (**H**) Immunostaining of phosphorated SMAD1/5/8 and ISL1 in the indicated cells treated with dox for 2 days. Three independent cultures were used for the experiment. Scale bar, 100 µm.

BE3 is located 66 kilobases downstream of the *BMP4* TSS, within the intronic region of a poorly characterized non-coding RNA gene (Fig. 6A). To determine whether BE3 functions as a distal enhancer for *BMP4* transcription *in situ*, we generated two independent ESC clones (ΔBE3#15 and #18) in which the genomic region corresponding to BE3 was deleted (Fig. S8A-B). These clones were engineered to express ELF3 in a dox-inducible manner, allowing *BMP4* transcription to be induced upon dox treatment (Fig. 6F). qPCR revealed that ELF3 induction led to robust *SPRR2A* upregulation in both parental and ΔBE3 clones (Fig. 6G), indicating that overall transcriptional responsiveness to ELF3 was preserved in ΔBE3 clones. However, *BMP4* expression was significantly reduced in ΔBE3 clones compared to parental cells, which was accompanied by decreased SMAD activation (Fig. 6G-H). Moreover, the expression of BMP-responsive TFs was markedly impaired in both ΔBE3 clones (Fig. 6H and Fig. S8C). These findings demonstrate that BE3 acts as a critical enhancer for BMP4 transcription in the ELF3-induction setting.

### Chromatin-level coordination of ELF3 with TEAD/TAZ in BMP4 distal enhancer

The coordinated action of transcription factors at enhancers of cell-type-specific genes often underlies the robust induction of these genes at specific time points during embryonic development. Using the BMP4 enhancer BE3 as a model, we examined whether such enhancer-level cooperation operates downstream of ELF3 induction. Motif analyses using HOMER identified three TEAD-binding motifs within the BE3 region, located adjacent to the putative ELF3-binding site (Fig. S9A). TEAD family TFs control transcription in concert with cofactors such as YAP and TAZ (*42*). TEAD1-4, YAP, and TAZ are all expressed and show nuclear localization (Fig. S9B-C). A luciferase reporter assay using a TEAD-responsive construct confirmed TEAD transcriptional activity in both ESCs and dox-treated cells (Fig. S9D). Importantly, a BE3 reporter lacking all TEAD-binding motifs failed to induce luciferase expression, highlighting the essential roles of these motifs in BE3 enhancer action (Fig. S9E-F). Moreover, co-transfection of TEADs with YAP or TAZ markedly increased BE3-driven reporter activity (Fig. S9G). These results suggest that the enhancer activity of BE3 can be potentiated by the TEAD/YAP/TAZ transcription module. However, despite these implications for a potential relationship between ELF3 and the TEAD/YAP/TAZ module at BE3, their co-expression did not result in a prominent synergistic enhancement of BE3-driven reporter activity (Fig. S9H). Importantly, ELF3 alone did not promote luciferase activity (Fig. S9H). These observations suggest that ELF3 may not act primarily as a transcriptional activator in this context but rather as a regulator of chromatin accessibility.

To address this speculation, we generated transgenic cells (tet-E3:ER^T2^-TAZ ESCs) that enable parallel manipulation of ELF3 and TAZ activities through dox- and 4-hydroxytamoxifen (4OHT)-responsive systems (Fig. 7A). In these cells, dox pre-treatment induces ELF3 expression, whereas 4OHT triggers nuclear translocation of the ER^T2^-TAZ fusion protein (Fig. 7A-B). Upon dox treatment, we observed a significant increase in *SPRR2A* mRNA, a known ELF3 target gene, whereas 4OHT administration led to upregulation of *CTGF*, a canonical TEAD/TAZ target gene (Fig. 7C) (*36, 43*). Interestingly, TAZ activation alone resulted in only faint induction of *BMP4* expression (Fig. 7C). Consistently, ChIP-qPCR analysis revealed minimal recruitment of ER^T2^-TAZ to the BE3 enhancer under these conditions (Fig. 7D). Given the limited accessibility of the BE3 region in ESCs, these results suggest that such a chromatin state may hinder TAZ recruitment to BE3, thereby constraining BMP4 transcription. In contrast, ELF3 pre-activation significantly increased TAZ binding to BE3 and robustly enhanced *BMP4* transcription (Fig. 7C-D). These findings indicate that ELF3 pre-activation is required for robust TEAD/TAZ recruitment to BE3 and efficient transcriptional activation of *BMP4*, indicating that ELF3 acts upstream of BMP4 induction primarily by increasing BE3 accessibility. Collectively, these findings position ELF3 as a regulator of enhancer accessibility that can enable engagement of transcriptional activators at specific loci, thereby facilitating transcriptional remodeling toward an amniotic-like cell state.

**Fig. 7.**
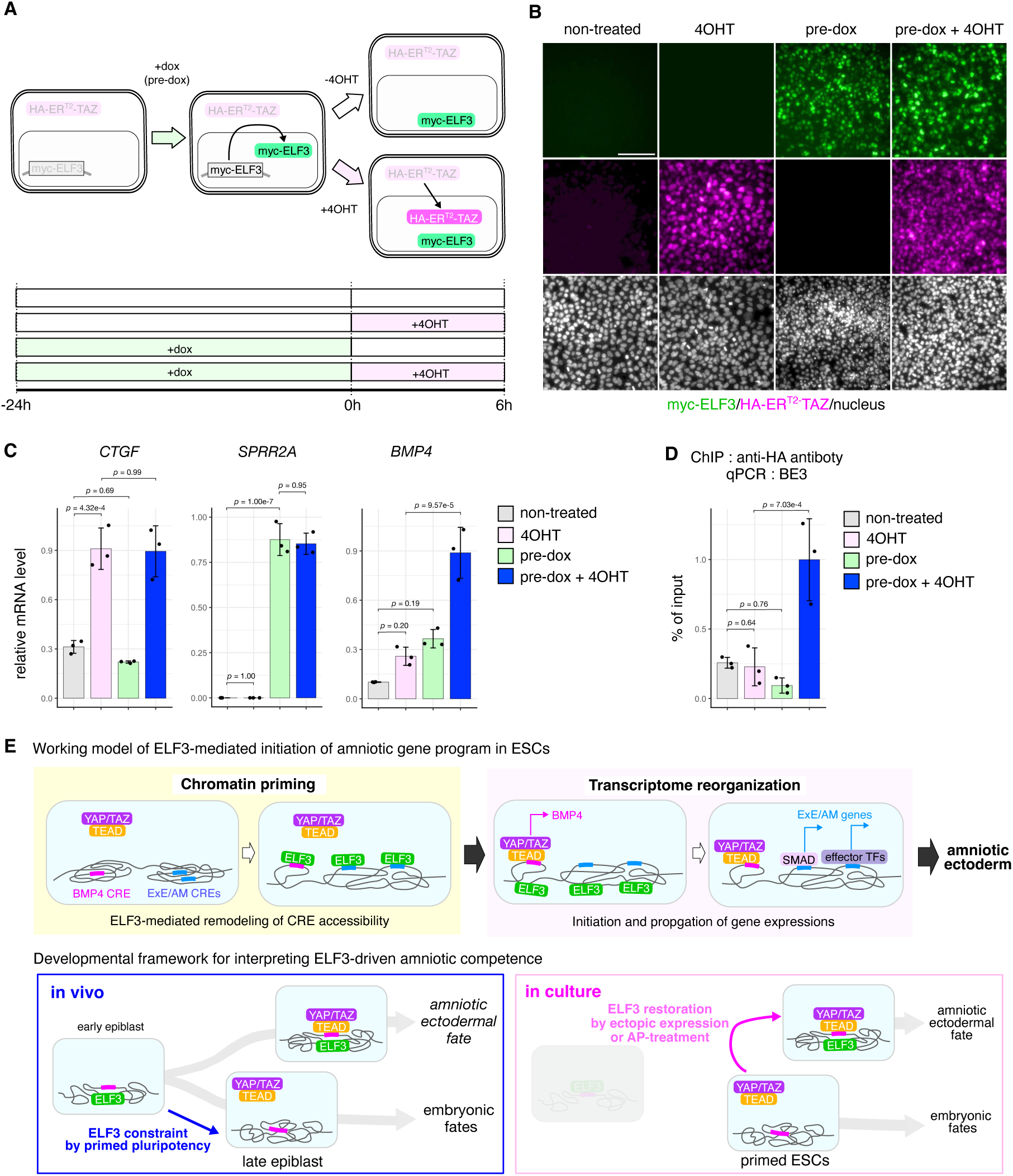
Chromatin priming role of ELF3 at the BE3 enhancer. (**A**) Schematic diagram illustrating the parallel manipulation for ELF3 and TAZ activities (top) and drug treatment timeline (bottom). (**B**) Immunostaining of HA-ER^T2^-TAZ and Myc-ELF3 in cells treated with 4-OHT, dox, both or untreated controls, as indicated in A. Scale bar, 100 µm. (**C**) qPCR quantification of *CTGF*, *SPRR2A*, and *BMP4* transcription in cells treated as indicated in A. Data are plotted as mean ± SD (n = 3 independent experiment). Statistical significance was determined by Tukey’s test. (**D**) ChIP-qPCR quantification of HA-ER^T2^-TAZ recruitment at the BE3 enhancer. Data are shown as mean ± SD (n = 3 independent experiments). Statistical significance was determined by Tukey’s test. (**E**) Summary diagram for ELF3-mediated activation of amniotic differentiation program. Working model (top). ELF3 increases accessibility at the BMP4 cis-regulatory element (BMP4 CRE) and other extraembryonic/amniotic CREs (ExE/AM CREs), enabling recruitment of additional transcriptional regulators (e.g., TEAD/YAP/TAZ) and activation of the downstream program (including BMP–SMAD-responsive programs and effector TFs). Developmental framework for interpreting ELF3-driven amniotic competence (bottom). ELF3 is constrained under primed pluripotency in implantation-stage embryos and in cultured PSCs. Restoring ELF3 action in culture (via transgene expression or AP treatment) re-enables access to latent amniotic competence.

## Discussion

Despite significant advances in understanding human development, the upstream mechanisms that initiate amniotic lineage segregation from the developing epiblast remain largely undefined (*44, 45*). Recent progress in PSC-based modeling of human embryogenesis provides an interesting possibility that amniotic differentiation could occur via cell-intrinsic mechanisms, which may resemble ancestral modes of amniotic lineage delamination, as seen for example in birds and reptiles. In this study, we identified ELF3 as a transcriptional entry factor that enables amniotic differentiation by facilitating the intrinsic activation of BMP signaling and subsequent activation of amniotic genes.

ELF3 is a member of the ETS family, a large group of TFs characterized by a conserved ETS DNA-binding domain (*46, 47*). ELF3 has been shown to play crucial roles in epithelial tissue formation and maintenance, vascular inflammation, and immune responses (*46–50*). Additionally, it has been reported to exert both tumor-promoting and -suppressive roles across multiple organs (*48–50*). Although ELF3 mediates these diverse responses by regulating context-dependent gene transcription, its role in early human development remains poorly explored. We found that ELF3 expression is low in primed ESCs but rapidly upregulated by a chemical perturbation of primed pluripotency (Fig. 3A-C), suggesting that the primed pluripotency-associated transcriptional circuit represses ELF3 expression. Interestingly, ectopic ELF3 expression in ESCs disrupts primed pluripotency and promotes transcriptional activation of the amniotic gene program. ELF3 deficiency had little effect on AP-induced pluripotency dissolution and GATA3^+^ ExEC differentiation (Fig. 3I and Fig. S5D), suggesting possible compensatory roles of other TFs, such as GATA6, IRX3, or related ETS factors. However, transcriptome analysis showed that AP-treated ELF3-deficient cells exhibited gene expression profiles less representative of the amnion-like transcriptional state (Fig. 3I-J). These results in loss- and gain-of-function experiments support a regulatory role of ELF3 in the amniotic gene program *in vitro*. Notably, our results suggest a mutually repressive relationship between ELF3 activity and the primed pluripotent state, which may tune the availability of amniotic competence.

To elucidate how ELF3 remodels the gene regulatory landscape, we performed ChIP-seq analyses, revealing that ELF3 binds genomic loci associated with extraembryonic phenotypes. ATAC-seq further demonstrated that many of these ELF3-bound regions exhibit limited accessibility in the primed pluripotent state but become accessible following ELF3 induction. Motif enrichment analysis revealed an overrepresentation of binding motifs for transcription factor families implicated in extraembryonic gene regulation, such as GATA, TEAD, and TFAP2, within ELF3-dependent accessible regions. A representative example was found at the BMP4 locus, where we identified a distal enhancer (BE3) as a direct ELF3 target (Fig. 6). Notably, BE3 contains multiple TEAD binding motifs adjacent to ELF3-binding sites (Fig. S9A). TEAD/YAP/TAZ are well established not only in regulating transcriptional programs but also in regulating characteristic behaviors of ESCs, including self-renewal and long-term survival (*51–55*). Using a parallel manipulation system, we demonstrated that TAZ alone was unable to bind to BE3 and drive BMP4 expression, whereas ELF3 expression enabled TAZ recruitment and robust *BMP4* transcription. These findings illustrate that ELF3 acts upstream of BMP4 transcriptional activation by increasing BE3 accessibility (Fig. 7E). We discuss this accessibility-enabling step in the context of “chromatin priming”, an early process in which increased cis-regulatory accessibility facilitates subsequent activation by additional regulatory inputs (*54, 55*).

Beyond *BMP4*, ELF3 also facilitates chromatin accessibility at several other amnion-related genes, including *GATA3*, *PRTG*, and *VGLL1* (Fig. 5E and Fig. S7E). Although we did not conduct detailed functional characterization of these loci, inhibitor experiments demonstrated that autocrine BMP-SMAD signaling contributes to the transcriptional induction of these genes (Fig. 1C), suggesting that ELF3 may play a broader role in rendering multiple amnion-related genes responsive to BMP-SMAD signaling. We therefore propose a working model for the intrinsic activation of an amniotic differentiation program, in which ELF3 first primes cis-regulatory elements of amnion genes by increasing their accessibility, and then their transcription is boosted by autocrine BMP–SMAD signaling (Fig. 7E). In this model, the cell-intrinsic induction of BMP4 itself may serve as a molecular bridge linking enhanced chromatin accessibility with the global transcriptomic reorganization.

While several questions remain unresolved, our findings provide important insights into extraembryonic lineage segregation during human development. A particularly intriguing question for future investigations is whether ELF3 participates in regulatory mechanisms underlying amniotic ectoderm formation *in vivo* during human embryogenesis. In addition, the *in vivo* spatiotemporal expression pattern of ELF3 provides further insight into its developmental roles: ELF3 is expressed from the morula stage to the early epiblast but downregulated along with epiblast maturation, persisting only in amniotic cells and trophoblasts (Fig. 1F). This expression pattern suggests that ELF3 is actively downregulated by primed pluripotency. From these findings, we speculate that low ELF3 levels and accompanying limited accessibility of extraembryonic gene regulatory elements in primed ESCs reflect the progressive restriction of extraembryonic differentiation competence during epiblast development (Fig. 7E). Consistent with these *in vivo* observations, ELF3 induction converts chromatin accessibility at amniotic gene loci from a low to a high state in primed ESCs. This suggests that epigenetic barriers that silence cis-regulatory elements of amniotic genes remain incompletely established at the primed stage; instead, amniotic competence is constrained by primed pluripotency-dependent repression of ELF3. These findings provide a mechanistic explanation for why primed PSCs remain capable of amnion-like differentiation and also prompt a reconsideration of their developmental potential. While traditionally viewed as a state poised for embryonic germ layer differentiation (*1, 56*), human primed pluripotency may also represent a transient window during which extraembryonic competence remains dormant but retrievable before its full closure.

This study offers important implications for the developmental regulation of differentiation competence. Further investigation into the regulatory functions of ELF3 and related ETS transcription factors in cell fate allocation will deepen our understanding of how extraembryonic differentiation potential is spatiotemporally defined during development. These insights may illuminate broader principles of transcription factor-mediated unveiling of latent differentiation competence, offering implications not only for lineage segregation mechanisms during human embryogenesis but also for advancing *in vitro* models of human development and future regenerative medicine.

## Materials and Methods

### Human pluripotent stem cells

Human ESC lines used in this study (KhES-1 and KthES13) were established previously in our institute with informed consent and provided to us in an anonymized form. The KhES-1-derived GATA3 reporter line (G3KI#2-5) was established as described in our previous study (*22*). The induced pluripotent stem cell line 253G1 was kindly provided by S. Yamanaka. All experiments using these cells were approved by the Ethics Committee of the Institute for Life and Medical Sciences, Kyoto University, and conducted in accordance with the guidelines and regulations of the Japanese government and the principles of the 2021 ISSCR Guidelines for Stem Cell Research. Although KhES-1 and G3KI#2-5 ESCs were mainly analyzed in this study, the other cell lines were also tested in some key experiments.

Undifferentiated human ESCs and iPSCs were cultured on feeder layers of mouse embryonic fibroblasts (MEF; Kitayama Labes; inactivated with 10 µg/ml mitomycin C and seeded at 4 × 10^5^ per 6 cm dish) in DMEM/F12/KSR medium (D-MEM/F12 (Sigma) supplemented with 20% KSR additive, 2 mM glutamine, 0.1 mM non-essential amino acids (Invitrogen), and 0.1 mM 2-mercaptoethanol). Recombinant human basic FGF (bFGF, 5 ng/ml, Wako) was added soon after seeding. For cell passaging, ESC colonies were detached by treating them with CTK dissociation solution at 37°C for 5-7 min, tapping the cultures, and then flushing them with a pipette, and recovering *en bloc* from the feeder layer (22). The detached ESC clumps were broken into smaller pieces by gently pipetting several times, and then these small clumps were transferred onto a MEF-seeded dish. For feeder-free cultures, contaminating MEF cells were removed by incubating the suspension on a gelatin-coated plate at 37°C for 2 h in the maintenance culture medium. In this procedure, MEF cells adhere to the dish’s bottom, but ESC clumps do not. The MEF-free ESC clumps were suspended in a MEF-conditioned medium and seeded onto a Matrigel substrate (BD Biosciences). The culture medium containing bFGF was refreshed daily until the next passage. When required, ESCs were cultured in medium containing 0.5 µg/ml dox, with daily media changes.

### Induction of GATA3^+^ ExEC Differentiation by AP treatment

ExEC differentiation was induced as previously described (*22, 57*). Human ESCs were seeded onto Matrigel-coated dishes, incubated overnight in a CO_2_ incubator, and then cultured in fresh medium containing A83-01 (1 µM) and PD173074 (0.1 µM). The medium was refreshed daily until analysis. For BMP signaling inhibition, cells were treated with AP in the presence of LDN-193189 (200 nM) and recombinant Chordin protein (0.5 µM).

### Plasmids, transfection, and generation of transgenic cells

The generation strategy transgenic cells in which transgene expression was induced by doxycycline supplementation, Piggybac (PB) transposon systems were employed as previously reported. HA-tagged cDNAs for candidate TFs were PCR-amplified from ESC genomic DNA and subcloned into pENTR1A Gateway entry vectors (Thermo Fisher Scientific). Following sequencing validation, cDNAs were transferred to PB transposon vectors (*pPB-hCMV*1-pA*) (*58*) by LR recombination reaction. For transient expression, cDNAs were similarly transferred into *pCAG-IRES-NeoR* or *pCAG-IRES-PuroR* vectors (*53, 57*). The *pCAG-hyPBase* expression vector was a gift from Dr. Yusa (*59*). The PB expression vector for HA-tagged ER^T2^-TAZ was constructed as previously described (*53*).

To generate polyclonal transgenic ESCs, PB vectors for candidate TFs were co-transfected with *pCAG-hyPBase* and *PB-CAG-rtTA-INeo* vectors using Lipofectamine Stem reagent (Invitrogen) (*22, 57*). A few days post-transfection, cells were replated onto G418-resistant MEF-coated dishes, and G418 selection (100 µg/ml) was initiated the following day. After weeks of selection, stable G418-resistant polyclonal pools were obtained. For g parallel manipulation of ELF3 and TAZ, four vectors (*PB-CAG-HA-ER^T2^-TAZ-INeo*, *PB-CMV*-Myc-ELF3*, *pCAG-hyPBase* and *PB-CAG-rtTA-INeo*) were co-transfected into ESCs. Two days later, cells were transferred onto puromycin-resistant MEF feeders and selected in medium containing 0.25 µg/ml puromycin for one week. Surviving cells were then transferred onto G418-resistant MEFs selected with 100 µg/ml G418 to obtain dual-resistant pools. Dox-dependent transgene induction and 4-OHT-dependent nuclear accumulation of HA-ER^T2^-TAZ were validated by immunostaining. When required, 4-OHT was added to the culture medium at 1 µg/ml.

### Luciferase reporter assays

To evaluate enhancer activity of ELF3-binding regions, corresponding genomic regions were PCR-amplified and subcloned upstream of the *NanoLuc* coding sequence in *pNL3.2[Nluc_minP]* vector (Promega) using an InFusion cloning system (Clontech). Deletion mutants were generated by inverse PCR and validated by sequencings. For TEAD reporters, 8×GTIIC sequences were PCR-amplified from the *8×GTIIC* luciferase vector (Addgene, #34615) and cloned into the *pNL3.2[Nluc_minP]* vector. Reporter plasmids were co-transfected with a firefly luciferase-expressing plasmid (*pGL3.2*, Promega) for normalization. Dual luciferase assays were performed using the Nano-Glo Dual-Luciferase Reporter Assay System (Promega), as previously described (*22*). In some experiments, TF expression plasmids were co-transfected. Luminescence was measured using a SpectraMax iD3 plate reader (Molecular Devices), and luciferase activity was represented as the NanoLuc-to-firefly luminescence ratio.

### Generation of mutant cell lines

The gene knockout by a Crispr-Cas9 genome editing system was done as previously described (*57*). Guide RNAs (gRNAs) targeting the ELF3 locus or the BE3 region were designed using a CRISPick webtool (https://portals.broadinstitute.org/gppx/crispick/public) (*60, 61*), as illustrated in Fig. S5C and S8A. The target sequences are listed in Supplementary Table S1. gRNA expression vectors were constructed using a *Cas9 sgRNA* backbone (Addgene, #68463), and *Cas9-2A-eGFP* expression plasmid (Addgene, #44719) was used for Cas9 expression. For ELF3 knockout, GATA3 reporter ESCs (G3KI#2-5) were co-transfected with gRNA and Cas9 vectors using Lipofectamine Stem reagent (Invitrogen). Two days later, eGFP-positive cells were isolated by flow cytometer (FACS Aria IIIu, BD Biosciences) and seeded at a low density (1,000 cells per 6-cm dishes) onto MEF-coated dished. Genome PCR and sequencing were used to identify biallelic deletions of targeted exons. Lack of ELF3 protein was confirmed by western blotting. BE3 deletion mutants were generated in KhES-1 cells following the same procedure. Two clones harboring deletion at the BE3 regions were identified by genomic PCR and sequencing. All primer sets were designed using a Primer3 and are listed in Supplementary Table S2.

### Quantitative real-time PCR

Total RNA was extracted using the RNeasy Mini Kit (Qiagen), and cDNA was synthesized using SuperScript II reverse transcriptase (Invitrogen). PCR reaction was prepared on 96-well plates using Power SYBR Green PCR Master Mix (Applied Biosystems), according to the manufacturer’s instructions. Primer sets were designed using a Primer-BLAST web tool (NCBI) and are listed in Supplementary Table S2. PCR reactions were run on a 7500 Fast Real-Time PCR System (Applied Biosystems). Transcript levels were quantified using a standard curve and normalized to GAPDH (*22, 53, 57*). Data were displayed either as arbitrary units or relative to control samples.

### Immunostaining analyses

Cells were fixed with 4% paraformaldehyde (PFA) at 4°C for 20 min, followed by permeabilization with 0.3% Triton-X100 solution. For VTCN1 and PRTG staining, cells were not permeabilized. After incubation in blocking buffer (e.g., 2% skim milk or 10 % normal donkey serum (Abcam) in 0.3% Triton-X100 solution), cells were incubated in the same buffer supplemented with primary antibodies. All primary antibodies used in this study are listed in Supplementary Table 3. Secondary antibodies conjugated with Alexa Fluor-488, -546, or -647 (Invitrogen) were used for visualization. Nuclei were counterstained with DAPI (WAKO). Images were acquired using a fluorescence microscope (LASX system, Zeiss) and processed using FIJI, an image processing package that bundles useful ImageJ plugins (ImageJ2 version 2.3.0). When required, image brightness and contrast were uniformly adjusted to enhance clarity.

### Western blotting

Cells were washed with PBS and lysed on the culture plate using IP lysis buffer (Thermo Fisher Scientific) for 10 min at 4 °C with gentle shaking. Whole cell lysates were harvested by pipetting. Immediately after adding the appropriate volume of 4 × LDS sample buffer (Invitrogen), samples were briefly sonicated for complete lysis and boiled. Protein extracts were resolved by SDS-PAGE and analyzed by western blotting as previously described^51^. A 5% nonfat dry milk solution was routinely used as a blocking reagent. Western blotting images were acquired using an Amersham Imager 600 (Fuji Film) and processed using FIJI. When required, image brightness and contrast were uniformly adjusted to improve clarity. All primary antibodies used in this study are listed in Table S3.

### Flow cytometry analyses

Flow cytometric analyses were performed using a flow cytometer (FACS Aria IIIu, BD Biosciences). Sample preparations for quantifying tdT fluorescence and detecting surface protein expression in live cells were performed as described before (*22, 57*). The primary antibody used for APA and VTCN1 staining is shown in Supplementary Table S3.

### Live imaging

For snapshots of live cells, cells were imaged by an inverted microscope (IX73, Olympus). For time-lapse live imaging, G3KI#2-5 ESCs were seeded onto a Matrigel-coated 35-mm µ-dish (Ibidi) (*22, 62*). On the following day, the culture medium was replaced with dox-containing medium, and time-lapse imaging was performed over 4 days at 10-minute intervals using an inverted microscope (IX81-ZDC, Olympus) equipped with a stepper filter wheel (Ludl) and a cooled EM-CCD camera (ImagEM, Hamamatsu Photonics). On day 2, the medium was replaced with a freshly prepared dox-containing medium.

### Bulk RNA-seq and data analyses

Total RNA was harvested as described above. Sequencing libraries were prepared from 1 μg total RNA using the NEBNext Ultra II Directional RNA Library Prep Kit (New England Biolabs) and sequenced by NextSeq550 (for dox-treated tetE3 clones) or NavaSeq6000 (for TF candidate screening and for E3KO characterization) (Illumina). Fastq files were generated using bcl2fastq (Illumina) and deposited in the Gene Expression Omnibus (GEO) database (GSE294600).

Sequence reads were aligned to the GRCh38 genome assembly using STAR (version 2.6.1d). Read counts and transcripts per million (TPM) values were calculated using RSEM (version 1.3.1). Read counts and TPMs were imported into the R platform (version 4.4.2) as the matrix data. Genes with total read counts < 10 across all given samples were excluded from further analyses. Differential gene expression was assessed using the edgeR (version 4.4.2) via exact tests. RNA-seq datasets obtained from the E3KO clones #21 and #28 were treated as biological replicates for statistical analysis. DEGs were defined as genes with *p* < 0.01, FDR < 0.05, and absolute fold change > 4. TPMs were used to assess relative transcript abundance. Principal component analyses were performed using the prcomp function. Gene set enrichment analysis was conducted using the fGSES (version 1.32.0). Previously published bulk RNA-seq data (GSE175977) were also used to analyze gene expression dynamics during AP-induced differentiation (*22*). A curated list of human transcription factors was obtained from HumanTFDB (http://bioinfo.life.hust.edu.cn/HumanTFDB#!/download) to extract TF from the given gene list (*63*).

### Single-cell RNA-seq data analyses

The scRNA-seq analyses of AP-induced differentiation (day 2–4) and its evaluation using the human embryo reference were done using Seurat (v5.4.0) as described in our previous report (*22, 64*), with some modifications. To examine ELF3, GATA6 and IRX3 expressions during early human development, we analyzed an integrated scRNA-seq dataset of human embryos using previously developed web-based tools, “human embryonic reference tool (version 1.1.1.17)” (*35*).

### ChIP experiments

Chromatin immunoprecipitation (ChIP) was performed using a Low Cell ChIP-seq Kit (Active Motif). A total of 50,000 cells (for H3K27me3, H3K27ac, H3K4me3, and p300) or 500,000 cells (for HA-ELF3, HA-ER^T2^-TAZ and MED1) were used for per reaction. Crosslinking and fixation were performed according to the manufacturer’s instructions. Sonication was performed using a Bioruptor sonication device (Sonic Bio) on 15 cycles (30 s-ON/30 s-OFF) in ice-cold water. Primary antibodies for ChIP are listed in Supplementary Table S3. Following de-crosslinking and purification, ChIP DNA quality was analyzed using High Sensitivity DNA Assay Kit on a Bioanalyzer (Agilent Technology). Recruitment of HA-ER^T2^-TAZ to the BE3 region was validated by ChIP-qPCR using an anti-HA antibody. Input DNA was prepared in parallel for normalization, and ChIP efficiency was calculated as a percentage of input. Primer sets used for detecting BE3 precipitation are listed in Supplementary Table S2.

### ChIP-seq and data analyses

Sequencing library preparation was performed by the Next Gen DNA Library Kit and Next Gen Indexing Kit, both included in the Low Cell ChIP-seq Kit (Active Motif). The libraries were sequenced bon a NextSeq550 (Illumina) to obtain paired-end reads. Fastq files were generated using bcl2fastq (Illumina) and deposited in the GEO database (GSE294602). Sequence reads were quality-checked with FastQC, trimmed using fastp, and aligned to the GRCh38 genome assembly using Bowtie2 with default parameters. PCR duplicates were removed using Picard (version 3.1.1). The aligned SAM files were converted to BigWig format using deeptools *bamCoverage* command with bins-per-million normalization (version 3.5.3), and visualized using Integrative Genomics Viewer (IGV, version 2.6.3, Broad Institute). For HA-ELF3 ChIP (dox-treated sample, day 2), peak calling was performed with MACS2 (version 2.2.7.2) using the parameter “-q 1e-10”, comparing ChIP samples to input controls. Peak distribution was analyzed using the deeptools *computeMatrix* command and visualized using *plotHeatmap* and *plotProfile* commands.

### ATAC experiments

ATAC experiments were performed following a previously described protocol (detailed protocol available at https://www.med.upenn.edu/kaestnerlab/protocols.html), using two biological replicates (*65*). A total of 50,000 cells were suspended in cold resuspension buffer (RSB; 10 mM Tris-HCl pH 7.5, 10 mM NaCl, and 3 mM MgCl2), centrifuged, resuspended in 50 μl of lysis solution (RSB with 0.1% NP40, 0.1% Tween-20, and 0.01% digitonin), incubated on ice for 3 min, and then diluted with 1 ml of wash buffer (RSB containing 0.1% Tween-20). The suspension was centrifuged to collect Nuclei. The transposition reaction mix consisted of 25 μl of 2×TD buffer, 2.5 μl of transposase (Nextera DNA Library Prep Kit, Illumina), 16.5 μl of PBS, 0.5 μl of 1% digitonin, 0.5 μl of 10% Tween-20, and 5 μl of water. Nuclei pellets were resuspended in 50 μl of transposition reaction mix and incubated at 37 °C for 30 min with shaking at 1,000 r.p.m. Transposed DNA was purified using the MinElute Reaction Cleanup Kit (Qiagen).

### ATAC-seq and data analyses

ATAC-seq library preparation was performed as described previously (*65*). Library quality was assessed using analyzed using a High Sensitivity DNA Assay Kit (Agilent Technology), and sequencing was performed on a NextSeq500 (Illumina) to generate single-end reads. The data were deposited in the GEO database (GSE294602). ATAC-seq reads were processed similarly to ChIP-seq data. Peak calling was performed using MACS2 with parameters “--nomodel --nolambda --shift -75 --extsize 150 -q 1e-10” without input control. Differentially accessible regions (DARs) between dox-treated and untreated samples were identified using DEseq2 via the DiffBind package (version 3.16.0). DARs were defined as regions with a *p* < 0.01, FDR < 0.05, and absolute fold-change > 2. Peak distribution was visualized using the deeptools as previously described. Overlapping peaks between ATAC-seq and ChIP-seq datasets were identified using bedtools *intersect* command. Motif Enrichment within peaks was assessed using the HOMER tool findMotifsGenome.pl (version 4.10) with parameters “-size 200 -mis 3 -p 6 -S 15”. Peak annotations were performed using the HOMER tool annotatePeaks.pl.

### Graphical presentation

Visualization of gene expression profiles in single-cell transcriptome data was performed using the *DimPlot*, *FeaturePlot*, *DotPlot* and *DoHeatmap* functions implemented in Seurat, with some modifications applied using ggplot2 (version 3.5.2). Flow cytometry data were visualized using FlowJo software (version 10.7.1). The visualization methods for ATAC-seq and ChIP-seq data were described above. All other graphs were generated using ggplot2.

## Statistical analysis

Data are presented as mean ± S.D., unless otherwise stated. The statistical tests used, exact n values, and exact P values are reported in the corresponding figures and figure legends. No statistical methods were used to predetermine sample size; sample sizes were chosen based on prior experience with the experimental system and published studies. Unless otherwise stated, each n represents an independent biological replicate (separate cultures or experiments). For qPCR, ChIP–qPCR, reporter assays, and flow cytometry, at least three independent experiments were performed. Western blotting and live imaging were performed at least twice. For immunofluorescence, at least three independent cultures were prepared, and key experiments were performed at least two times. For comparisons between two groups, two-sided unpaired Student’s t-tests were used. For comparisons among multiple groups, one-way ANOVA followed by Tukey’s post hoc test (all pairwise comparisons) or Dunnett’s post hoc test (versus control) was performed. Analyses were performed in R. Statistical significance was defined as *p* < 0.05 unless otherwise stated (TF screening experiments used *p* < 0.001). Bulk RNA-seq and bulk ATAC-seq were performed with two independent biological replicates per condition/time point. ChIP-seq was performed once per condition and used for qualitative analyses without replicate-based statistical testing. Differential expression analysis of bulk RNA-seq data was performed using edgeR, with *p*-values adjusted for multiple testing using the Benjamini–Hochberg false discovery rate (FDR). Differential accessibility analysis of bulk ATAC-seq data was performed using DiffBind with DESeq2, with *p*-values adjusted using FDR.

## Acknowledgments

We are grateful to the Liaison Laboratory Research Promotion Center (LILA) at IMEG Kumamoto University and Single-cell Genome Information Analysis Core (SignAC) at WPI-ASHBi Kyoto University for sequencing; Y. Mii, R. Tsutsumi, and Z. Whang for discussions and fruitful comments on the manuscript; A. Ohgushi and all members of our laboratory for support and encouragement. This work was supported by JSPS KAKENHI Grant Number 23K23795 and 24K22035 (to M.O.), Joint Usage/Research Center for Developmental Medicine IMEG Kumamoto University (to M.O.), and was partially supported by JSPS KAKENHI Grant Number 23H04933 (to M.E.) and MEXT Promotion of Development of a Joint Usage/Research System Project Grant Number JPMXP1323015486 (to M.E.).

## Author contributions statement

M.O. designed the project, conducted the experiments and data analyses, and wrote the manuscript; K.H. conducted some experiments; H.N. and M.E. supported the data analyses. All authors reviewed the manuscript.

## Data availability

The authors declare that all data supporting the findings of this study are included within the article and its Supplementary Information or from the corresponding author upon reasonable request. Row data of RNA-seq, ChIP-seq and ATAC-seq are available from the Gene Expression Omnibus (GSE294600, GSE294602).

