## Supplemental Figures for "Chromatin priming by transcription factor ELF3 confers transcriptional competence for amniotic differentiation on primed human embryonic stem cells"

Supplementary Figures

Figure S1. Expression profile of lineage markers during AP-induced differentiation.
Figure S2. Expression screening for TFs capable of inducing GATA3+ ExECs.
Figure S3. Comprehensive characterization of ELF3-induced ExECs.
Figure S4. ELF3-induced ExECs from different PSC lines.
Figure S5. Characterization of ELF3 knockout ESCs.
Figure S6. Profiling of ELF3 chromatin bindings.
Figure S7. Chromatin accessibility changes upon ELF3 induction.
Figure S8. Enhancer validation of ELF3-bound elements around the BMP4 gene.
Figure S9. TEAD/YAP/TAZ regulation of BE3 activity.

Supplementary Tables

Table S1. Target sequences for gRNAs.
Table S2. Primer sequences.
Table S3. Antibody list.

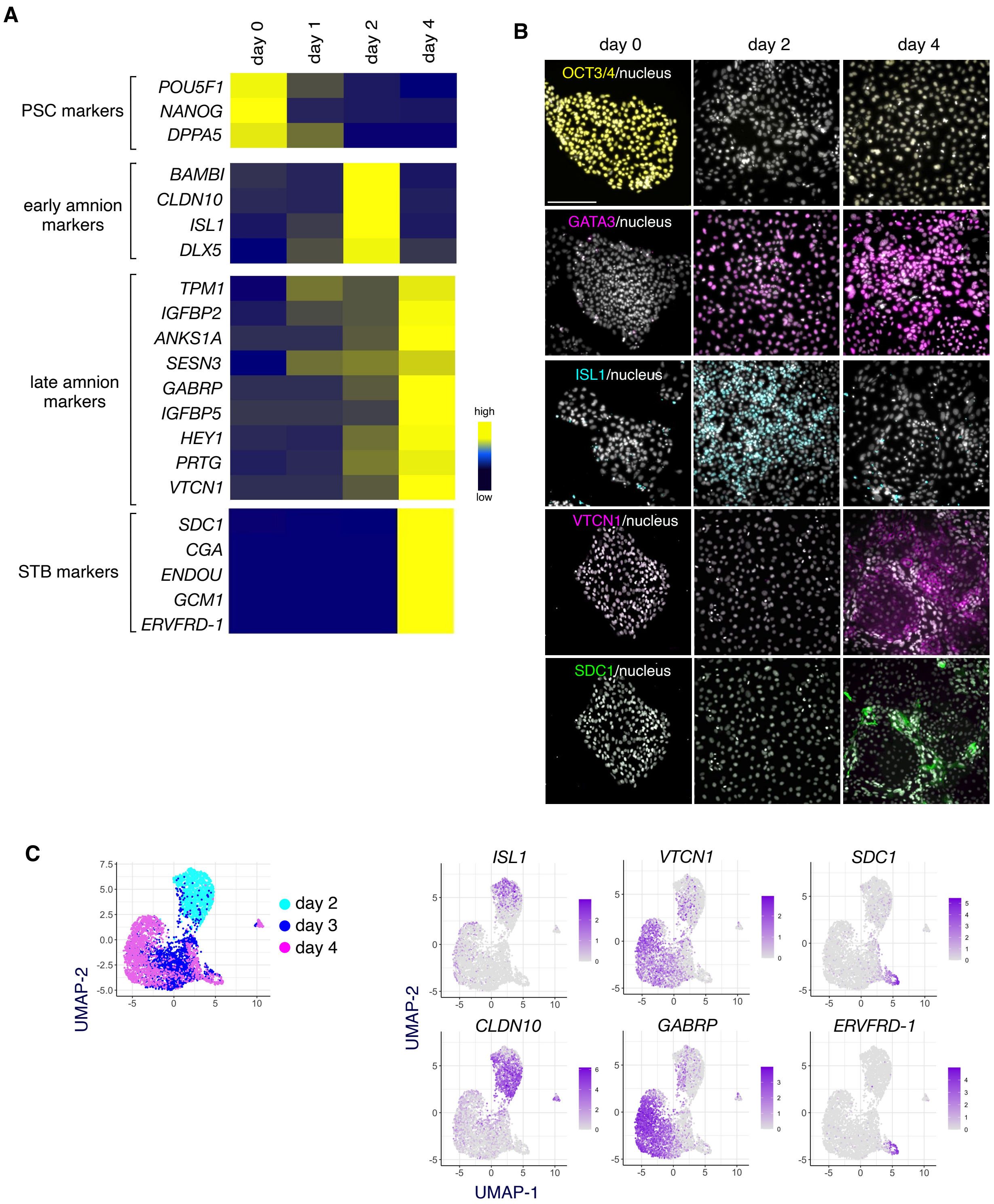


**Figure S1. Expression profile of lineage markers during AP-induced differentiation. A**, Heatmap showing expression dynamics of markers for PSCs, early and late amnion, and STBs during AP-induced differentiation. Amnion markers include genes proposed in previous studies (Rostovskaya et al., Cell Stem Cell, 2022; Sekulovski et al., elife, 2024; Ohgushi et al., Cell Reports, 2022). Gene expression is shown as a scaled TPM (mean of two replicates). **B**, Immunostaining of OCT3/4, GATA3, ISL1, VTCN1, and SDC1 at the indicated time points of AP-induced differentiation. Two independent experiments were done. Scale bar, 200 µm. **C**, Re-anaylyses of our privious scRNA-seq data. Expression matrics of scRNA-seq experiments on AP-treated cells (day 2, 3, and 4; Ohgushi et al., Cell Reports, 2022) were merged and projected onto the same UMAP. Expression profiles of individual lineage markers (ISL1/CLDN10 for early amnion, VTCN1/GABRP for late amnion, and SDC1/ERVFRD-1 for STB) were shown on the UMAP.

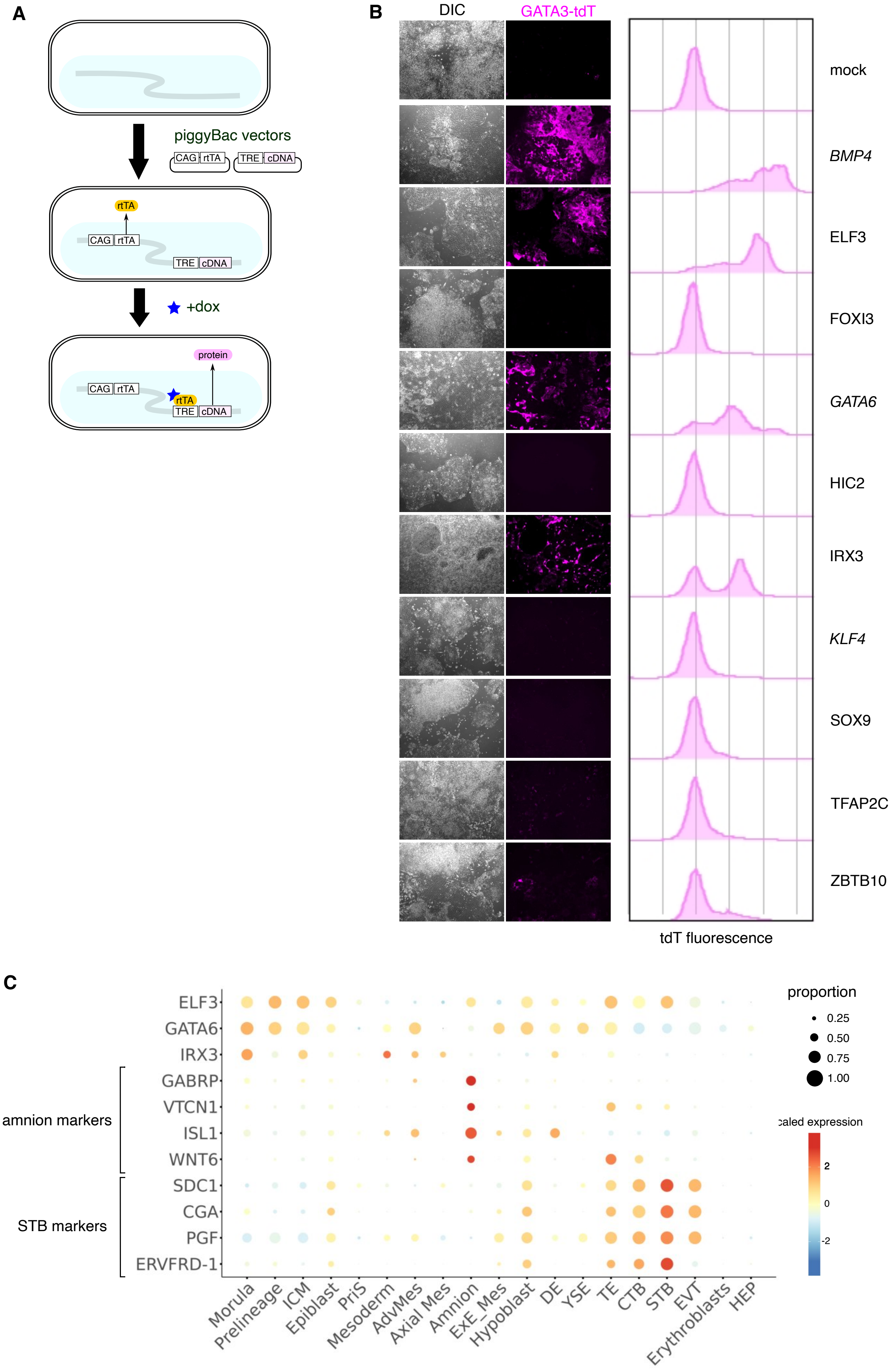
A
piggyBac vectors
CAG rtTA
TRE cDNA
rtTA
+dox
protein
B
DIC
GATA3-tdT
mock
BMP4
ELF3
FOXI3
GATA6
HIC2
IRX3
KLF4
SOX9
TFAP2C
ZBTB10
tdT fluorescence
C
ELF3
GATA6
IRX3
GABRP
VTCN1
ISL1
WNT6
SDC1
CGA
PGF
ERVFRD-1
amnion markers
STB markers
Morula
Prelineage
ICM
Epiblast
PriS
Mesoderm
AdvMes
Axial Mes
Amnion
EXE_Mes
Hypoblast
DE
YSE
TE
CTB
STB
EVT
Erythroblasts
HEP
proportion
0.25
0.50
0.75
1.00
scaled expression
2
0
-2

**Figure S2. Expression screening for TFs capable of inducing GATA3+ ExECs. A,** Schematic diagram for dox-inducible TF expression system. **B**, Transgenic cells engineered to induce individual TFs in a dox-dependent manner were treated with dox for 4 days. Representative images of phase-contrast and GATA3-tdTomato fluorescence images of each transgenic cell line are shown alongside corresponding flow cytometry panels (3 independent experiments). **C**, Dot plot showing cross-lineage comparison of *ELF3*, *IRX3*, and *GATA6* mRNA expressions in the human embryo scRNA-seq dataset (Tyser et al., 2021, Xiang et al., 2020). Representative genes for the amnion and STB were also shown.

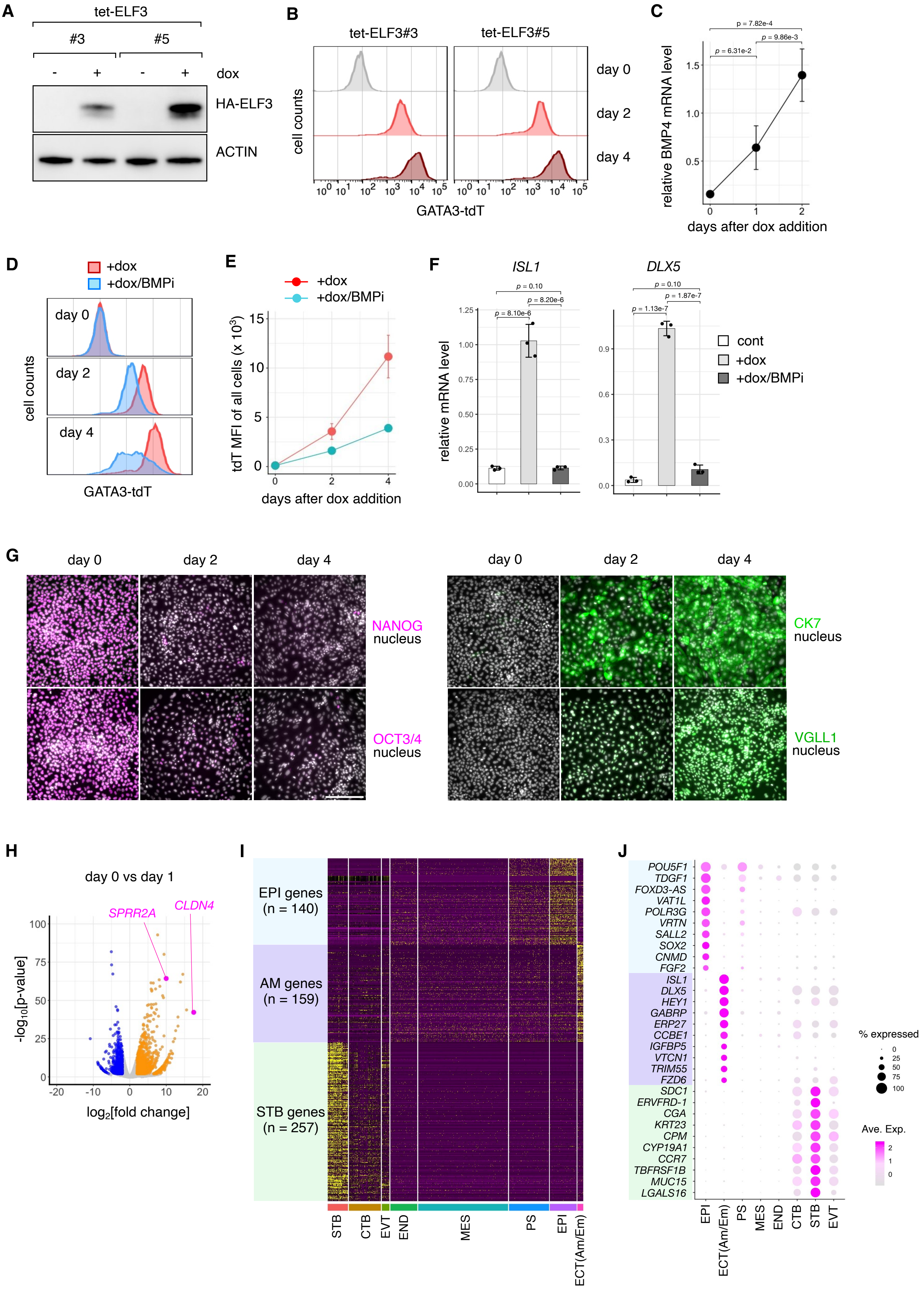
A
tet-ELF3
#3
#5
dox
HA-ELF3
ACTIN
B
tet-ELF3#3
tet-ELF3#5
day 0
day 2
day 4
cell counts
GATA3-tdT
C
relative BMP4 mRNA level
days after dox addition
p = 7.82e-4
p = 6.31e-2
p = 9.86e-3
D
+dox
+dox/BMPi
cell counts
GATA3-tdT
E
tdT MFI of all cells (x 10³)
days after dox addition
F
ISL1
DLX5
relative mRNA level
cont
+dox
+dox/BMPi
G
day 0
day 2
day 4
NANOG
nucleus
OCT3/4
nucleus
CK7
nucleus
VGLL1
nucleus
H
day 0 vs day 1
SPRR2A
CLDN4
-log10[p-value]
log2[fold change]
I
EPI genes (n = 140)
AM genes (n = 159)
STB genes (n = 257)
STB
CTB
EVT
END
MES
PS
EPI
ECT(Am/Em)
J
POU5F1
TDGF1
FOXD3-AS
VAT1L
POLR3G
VRTN
SALL2
SOX2
CNMD
FGF2
ISL1
DLX5
HEY1
GABRP
ERP27
CCBE1
IGFBP5
VTCN1
TRIM55
FZD6
SDC1
ERVFRD-1
CGA
KRT23
CPM
CYP19A1
CCR7
TBFRSF1B
MUC15
LGALS16
% expressed
Ave. Exp.

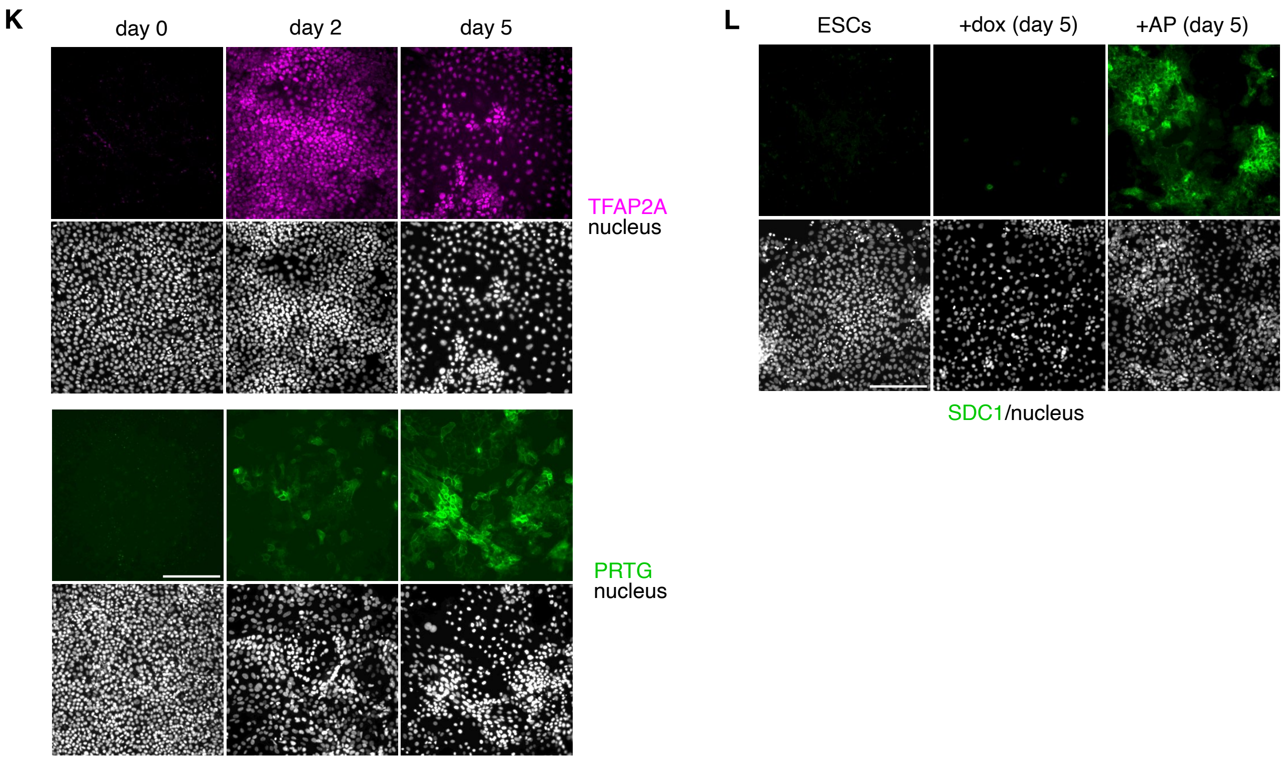


**Figure S3. Comprehensive characterization of ELF3-induced ExECs. A**, Representative WB images showing HA-ELF3 induction in tet-ELF3 clones (#3 and #5) from two independent experiments. Cells were treated or untreated with dox for 2 days. Actin was used as a loading control. **B**, Flow cytometry panels showing GATA3-tdTomato expression kinetics of dox-treated (2 or 4 days) and untreated clones. Three experiments were done. **C**, qPCR quantification of *BMP4* transcription after dox treatment. Data are plotted as mean ± SD (n = 3 independent experiments). Statistical significance was determined by Tukey's test. **D-E**, Flow cytometry panels (d) and graph of mean fluorescence intensity (MFI) of all cells (e) at the indicated time points following dox treatment (2 or 4 days) in the presence (red) or absence (light blue) of BMP inhibitors (BMPi). Three experiments were done. **F**, qPCR quantification of *ISL1* and *DLX5* levels in cells treated with dox with or without of BMPi for 2 days. Data are shown as mean ± SD (n = 3 independent experiments). Statistical significance was determined by Tukey's test. **G**, Immunostaining of pluripotency-associated markers (OCT3/4, NANAG) and extraembryonic markers (CK7, VGLL1) at the indicated time points after dox supplementation. Three independent cultures were used for the experiment. Scale bar, 100 µm. **H**, Volcano plot comparing dox-treated cells (for 24 h) versus control ESCs (two biological replicates per condition). Upregulated genes (orange) and downregulated (blue) genes are highlighted, respectively. ELF3 targets *SPRR2A* and *CLDN4* are marked in pink. **I-J**, Expression profile of lineage marker genes across embryonic clusters using the integrated human embryo scRNA-seq dataset. Heatmap showing expression of epiblast (140), amnion (159) and STB (257) markers (i). Dot plot showing expression of representative genes from each lineag (j). **K**, Immunostaining of the indicated amnion markers at the indicated time points after dox-mediated ELF3 induction. Three independent cultures were used for the experiment. Scale bar, 100 µm. **L**, Immunostaining of the SDT marker SDC1 at 5 days after dox-nediated ELF3 induction. Immunostaining images of AP-treated cells at day 5 were presented as posivive controls. Three independent cultures were used for the experiment. Scale bar, 100 µm.

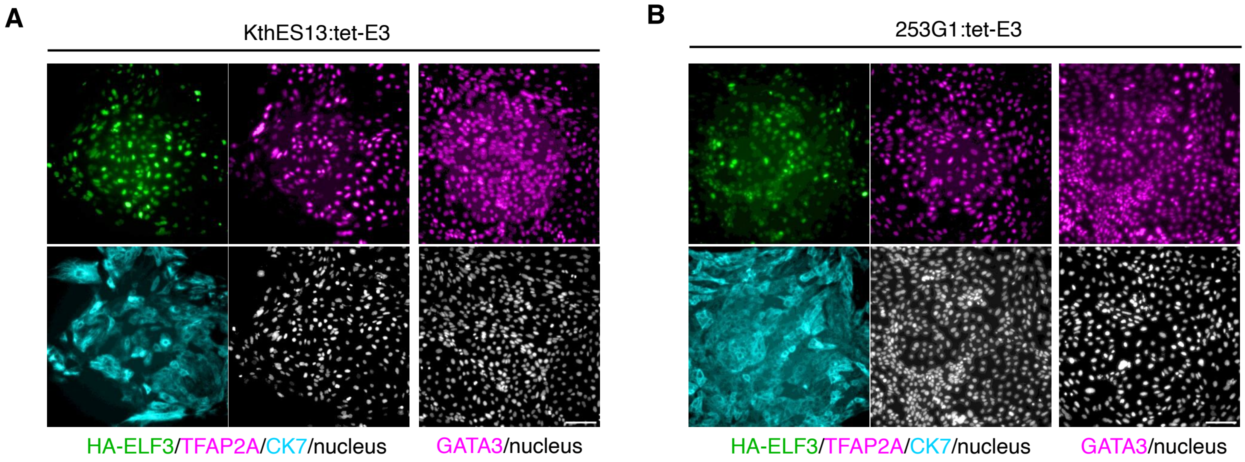


**Figure S4. ELF3-induced ExECs from different PSC lines. A-B**, KthES13 (a human ESC line; a), and 253G1 (a human induced pluripotent stem cell line; b), were engineered to express HA-ELF3 upon dox supplementation (as in Supplementary Fig. 2a). These cells were treated with dox for 5 days, and the expression of extraembryonic markers (TFAP2A, CK7, GATA3) were confirmed by immunostaining. Two independent cultures were used for the experiment. Scale bars; 100 µm.

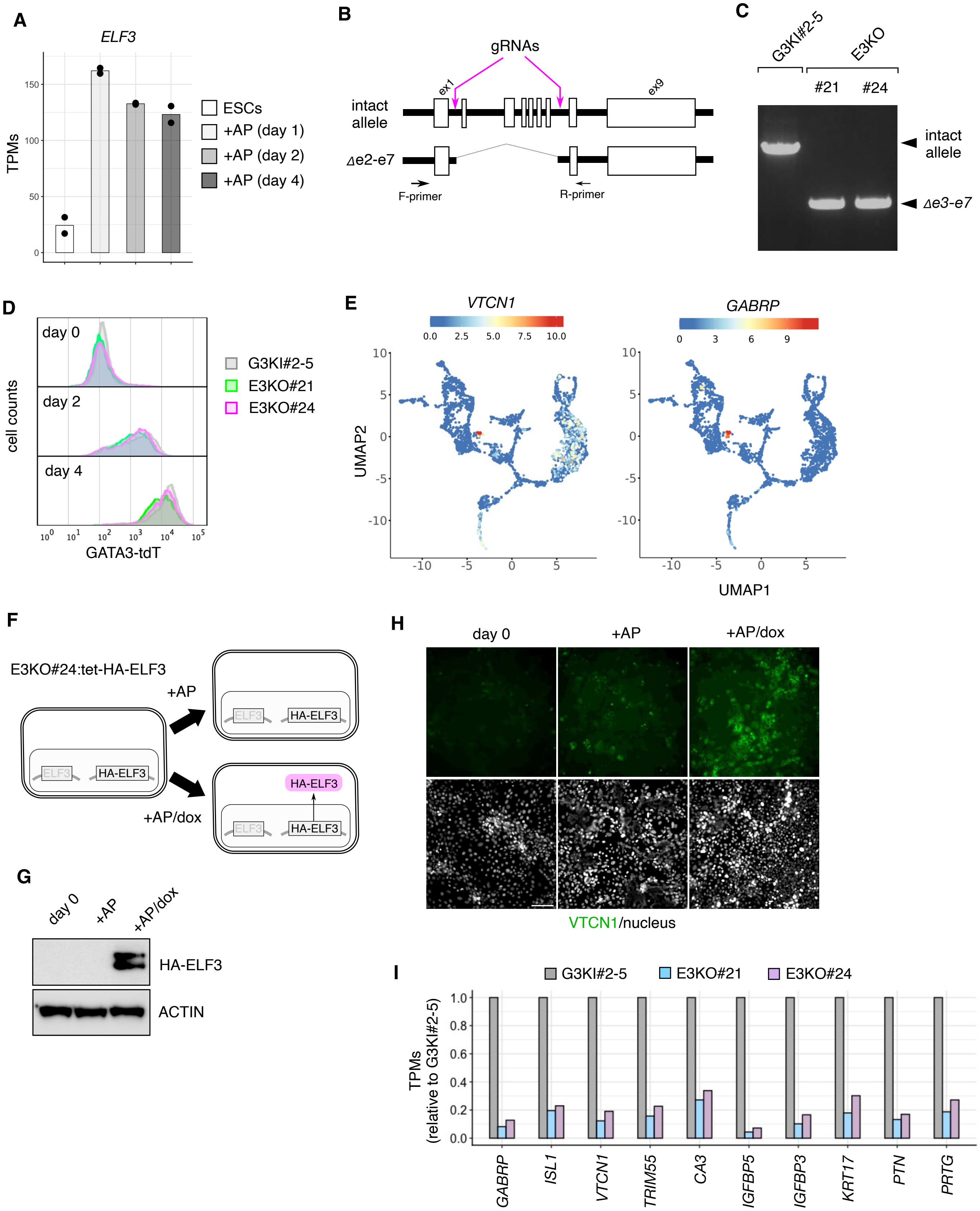

A
ELF3
TPMs
ESCs
+AP (day 1)
+AP (day 2)
+AP (day 4)
B
gRNAs
intact allele
Δe2-e7
ex1
ex9
F-primer
R-primer
C
G3KI#2-5
E3KO
#21
#24
intact allele
Δe3-e7
D
cell counts
day 0
day 2
day 4
GATA3-tdT
G3KI#2-5
E3KO#21
E3KO#24
E
VTCN1
GABRP
UMAP2
UMAP1
F
E3KO#24:tet-HA-ELF3
+AP
+AP/dox
ELF3
HA-ELF3
G
day 0
+AP
+AP/dox
HA-ELF3
ACTIN
H
day 0
+AP
+AP/dox
VTCN1/nucleus
I
G3KI#2-5
E3KO#21
E3KO#24
TPMs (relative to G3KI#2-5)
GABRP
ISL1
VTCN1
TRIM55
CA3
IGFBP5
IGFBP3
KRT17
PTN
PRTG

**Figure S5. Characterization of ELF3 knockout ESCs. A,**Time-course quantification of *ELF3* transcription during AP-nduced differentiation by RNA-seq. Data are shown as mean TPM values (two replicates). **B**, Schematic of ELF3 gene disruption strategy. Two gRNAs were designed to target intron 1 and 7 of the ELF3 gene. F- and R- primers are used for genomic PCR to confirm deletion of exon 2-7 (indicated as Δe2-e7). **C**, PCR confirmation using genomic DNA from candidate clones (#21, #28) and parental G3KI#2-5 ESCs. Representative images were shown from two independent experiments. **D**, Flow cytometry panels showing GATA3-tdTomato expression kinetics. ELF3 knockout clones #21 (green), #24 (magenta), and the parental (gray) were untreated or treated with AP for 0, 2 or 4 days. **E,** Expression patterns of *VTCN1* and *GABRP* mRNA on the embryo reference UMAP. Lineage annotations were shown in Figure 1f. **F**, Schematic diagram for ELF3 add-back experiments. E3KO#24 clone was engineered to express HA-ELF3 in response to dox supplementation (E3KO#24:tet-HA-ELF3). **G**, WB analyses showing HA-ELF3 induction in E3KO#24:tet-HA-ELF3 cells treated with AP only and AP with dox (for 5 days), or in the untreated. Representative images were shown from two experiments. Actin was used as a loading control. **H,** Immunostaining of VTCN1 in E3KO#24:tet-HA-ELF3 cells treated as in i. Three independent cultures were used for the experiment. Scale bar, 100 µm. **I**, Expression levels of representative amnion-related genes in ELF3 knockout clones (#21, #28; light blue and pink, respectively) and parental G3KI#2-5 (gray) after 4 days of AP treatment. Data are shown as relative TPM values to the control.

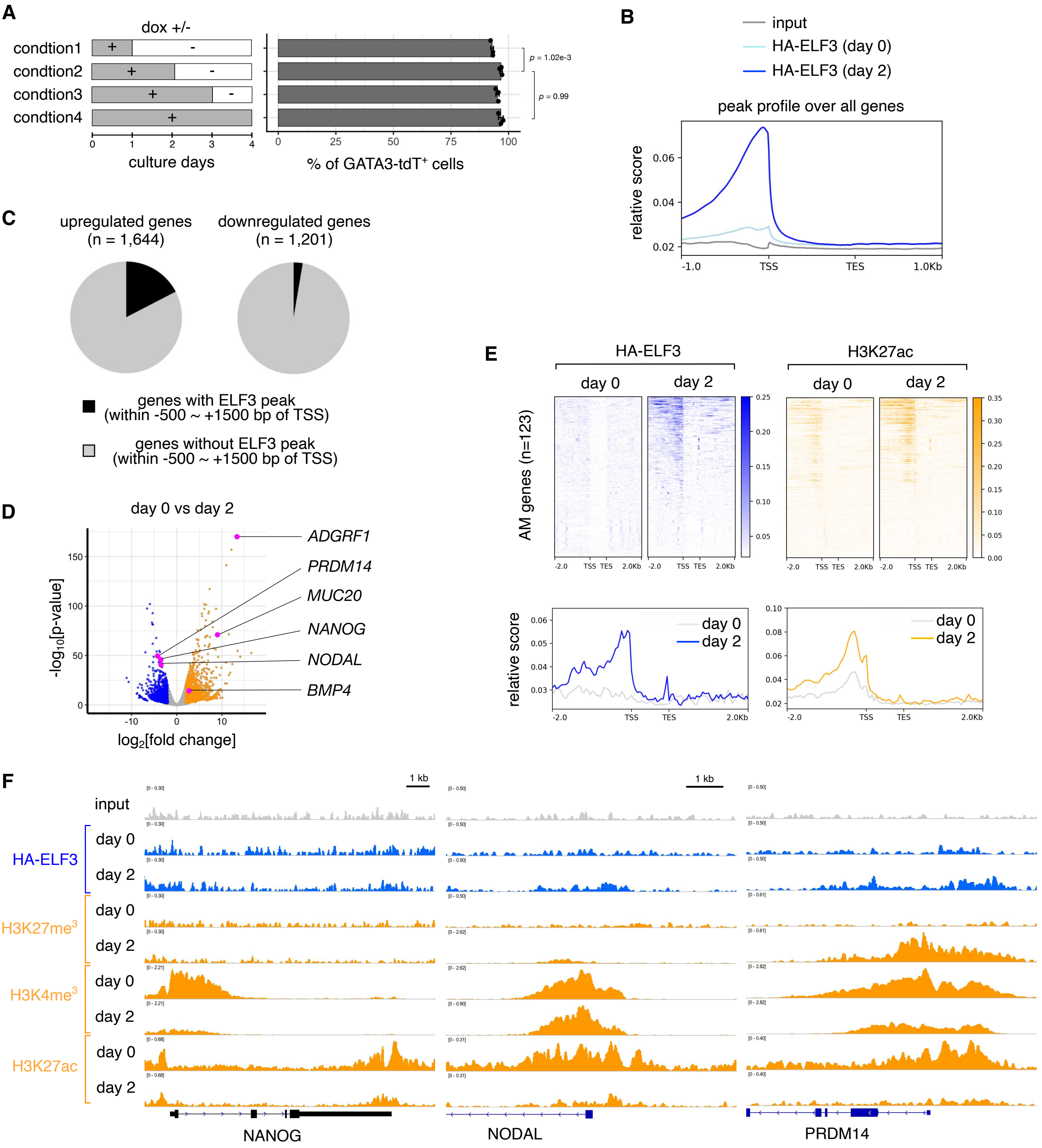


**Figure S6. Profiling of ELF3 chromatin bindings. A**, Time window analysis of dox treatment for generating GATA3+ ExECs. Dox was present during the periods indicated as gray bars. Flow cytometry was performed at day 4. Three experiments were done. **B**, Coverage profiles of HA-ELF3 over all genes in dox-treated (2 days) and untreated cells. ChIP peaks of isotype-matched control IgG were shown as input. **C**, Percentages of upregulated or downregulated genes with ELF3 binding within 1,500 bp upstream and 500 bp downstream of the TSS. **D**, Volcano plot of dox-treated cells (2 days) versus control ESCs. Upregulated and downregulated genes are shown as orange and blue dots, respectively. Selected example genes for genome track views are indicated in pink dots. **E**, HA–ELF3 occupancy and H3K27ac enrichment across amnion-associated gene loci before (day 0) and after induction (day 2). Genes are ranked by the mean HA–ELF3 signal at day 2. The amnion-associated gene list is the same in Fig. 2G. **F**, Genomic deposition of normalized ChIP signals at pluripotency-associated genes.

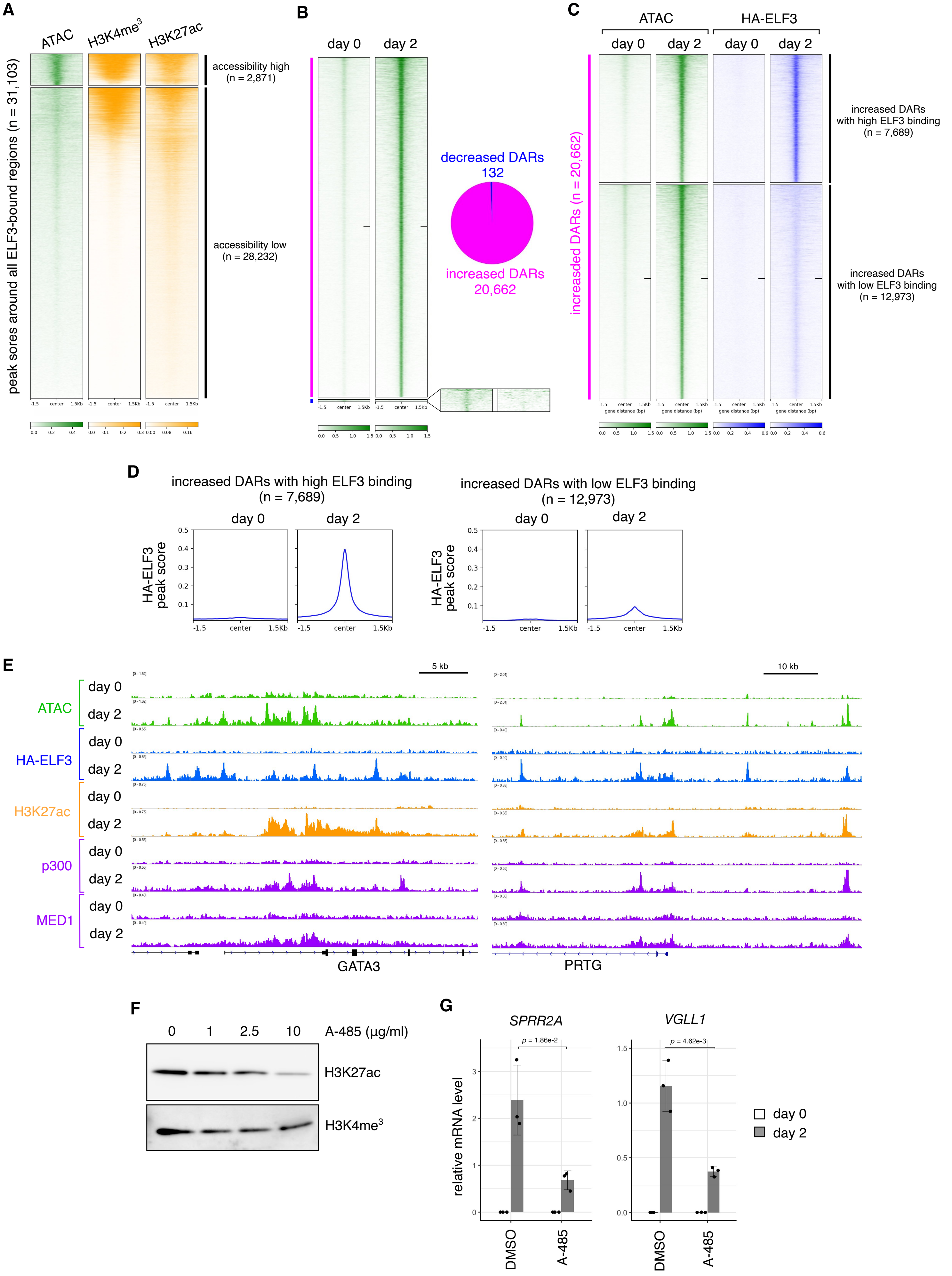
A
ATAC
H3K4me3
H3K27ac
peak sores around all ELF3-bound regions (n = 31,103)
accessibility high (n = 2,871)
accessibility low (n = 28,232)
B
day 0
day 2
decreased DARs 132
increased DARs 20,662
C
ATAC
HA-ELF3
day 0
day 2
increased DARs (n = 20,662)
increased DARs with high ELF3 binding (n = 7,689)
increased DARs with low ELF3 binding (n = 12,973)
gene distance (bp)
D
increased DARs with high ELF3 binding (n = 7,689)
increased DARs with low ELF3 binding (n = 12,973)
HA-ELF3 peak score
-1.5
center
1.5Kb
E
5 kb
10 kb
ATAC
HA-ELF3
H3K27ac
p300
MED1
GATA3
PRTG
F
0
1
2.5
10
A-485 (µg/ml)
H3K27ac
H3K4me3
G
SPRR2A
VGLL1
p = 1.86e-2
p = 4.62e-3
relative mRNA level
DMSO
A-485
day 0
day 2

**Figure S7. Chromatin accessibility changes upon ELF3 induction. A**, Coverage heatmaps showing chromatin accessibility and the indicated histone marks at ELF3 binding loci (same as indicated in Fig. 4a) in ESCs. Profiles are separated into low and high accessibility, determined by the presence or absence of ATAC peaks. **B**, Coverage heatmaps showing differentially accessible regions (DARs) in dox-treated (2 days) or untreated cells. An enlarged view of decreased DARs is also shown. The pie chart indicates the ratio of increased to decreased DARs. **C-D**, Coverage heatmap (c) and profile (d) of increased DARs and ELF3 binding. Profiles are separated into low and high ELF3 binding groups (as determined by DESeq2 using DiffBind). **E**, Genomic deposition of normalized ATAC and ChIP signals at amnion-related genes (GATA3 and PTRG). **F**, Representative WB images confirming inhibitory effects of A-485 (p300/CBP inhibitor) on global H3K27me3 levels in ESCs (two independent experiments). H3K4me3 was examined as a negative control. Cells were treated with dox in the presence or absence of A-485 at different concentrations for 2 days. **G**, qPCR quantification of *SPRR2A* and *VGLL1* transcription. Increased p300 deposition was observed at the cis-elements of these genes (see Fig. 5E). Cells were untreated or treated with dox in the presence or absence of A-485 (10 μM) for 2 days. Data are plotted as means ± SD (n = 3 independent experiments). Statistical significance was determined by two-sided Student' s t-test.

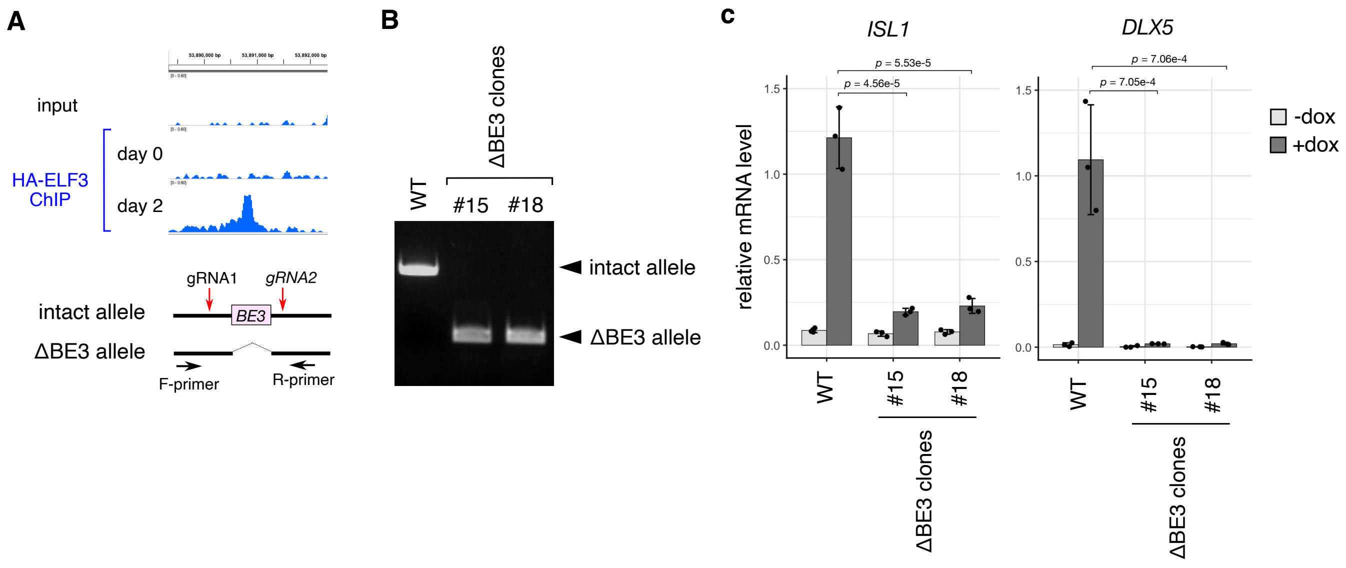


**Figure S8. Enhancer validation of ELF3-bound elements around the BMP4 gene. A**, Schematic diagram for the genetic disruption of BE3. Two gRNAs were designed to flank the genomic region containing the BE3 sequence. F- and R- primers are used for genomic PCR to confirm deletion (indicated as ΔBE3). **B**, PCR confirmation using genomic DNA extracted from candidate clones (#15, #18) and parental KhES-1 cells. Representative images were shown from two independent experiments. **C**, qPCR quantification of *ISL1* and *DLX5* transcription in the mutant clones and parental cells. Cells were untreated or treated with dox for 2 days. Data are shown as means ± SD (n = 3 independent experiments). Statistical significance was determined by Dunnett's test (vs dox-treated parental cells).

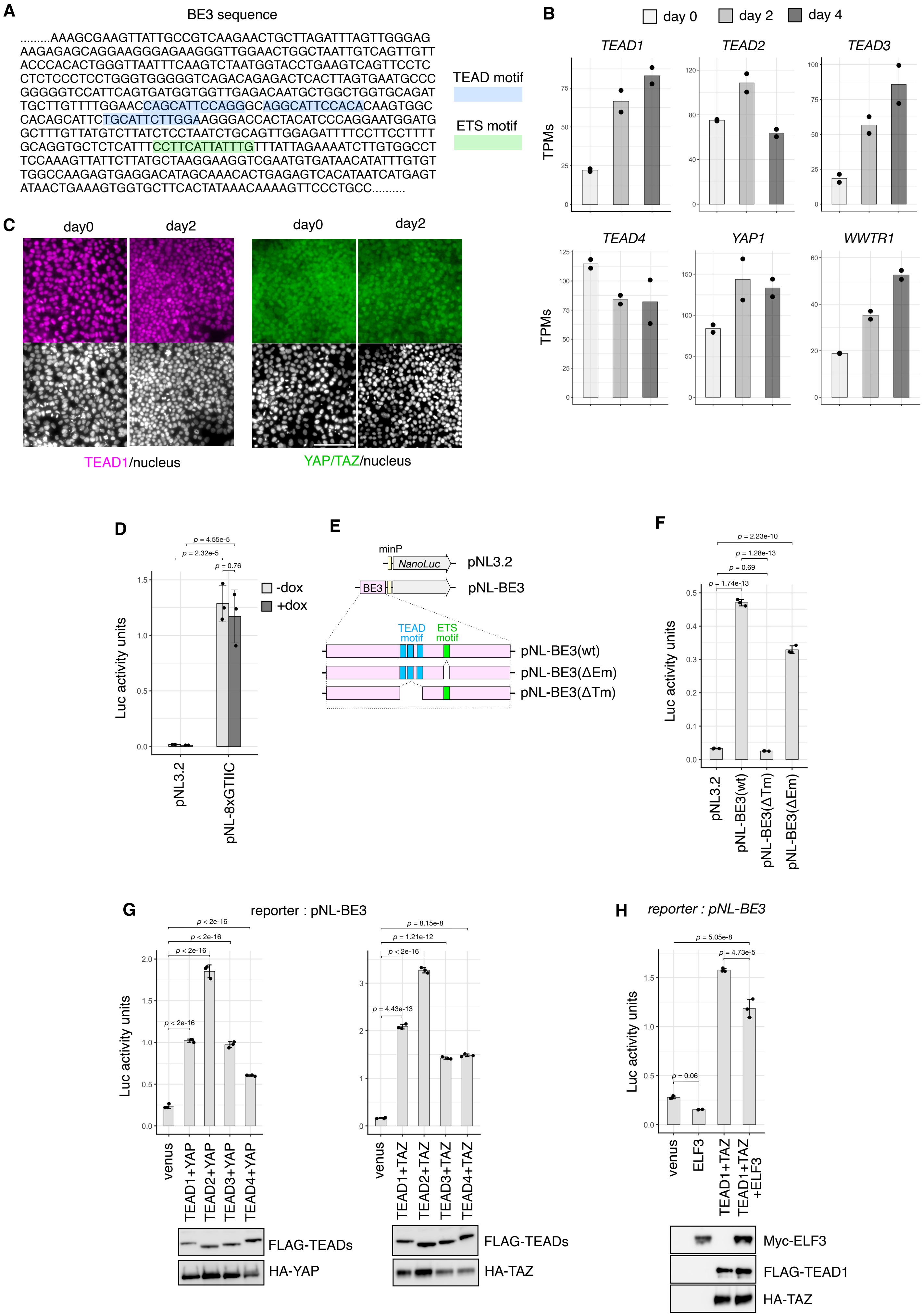
A
BE3 sequence
.........AAAGCGAAGTTATTGCCGTCAAGAACTGCTTAGATTTAGTTGGGAG
AAGAGAGCAGGAAGGGAGAAGGGTTGGAACTGGCTAATTGTCAGTTGTT
ACCCACACTGGGTTAATTTCAAGTCTAATGGTACCTGAAGTCAGTTCCTC
CTCTCCCTCCTGGGTGGGGGTCAGACAGAGACTCACTTAGTGAATGCCC
GGGGGTCCATTCAGTGATGGTGGTTGAGACAATGCTGGCTGGTGCAGAT
TGCTTGTTTTGGAACCAGCATTCCAGGGCAGGCATTCCACACAAGTGGC
CACAGCATTCTGCATTCTTGGAAGGGACCACTACATCCCAGGAATGGATG
GCTTTGTTATGTCTTATCTCCTAATCTGCAGTTGGAGATTTTCCTTCCTTTT
GCAGGTGCTCTCATTTCCTTCATTATTTGTTTATTAGAAAATCTTGTGGCCT
TCCAAAGTTATTCTTATGCTAAGGAAGGTCGAATGTGATAACATATTTGTGT
TGGCCAAGAGTGAGGACATAGCAAACACTGAGAGTCACATAATCATGAGT
ATAACTGAAAGTGGTGCTTCACTATAAACAAAAGTTCCCTGCC..........
TEAD motif
ETS motif
B
day 0
day 2
day 4
TEAD1
TEAD2
TEAD3
TEAD4
YAP1
WWTR1
TPMs
C
day0
day2
day0
day2
TEAD1/nucleus
YAP/TAZ/nucleus
D
p = 4.55e-5
p = 2.32e-5
p = 0.76
-dox
+dox
Luc activity units
pNL3.2
pNL-8xGTIIC
E
minP
NanoLuc
pNL3.2
BE3
pNL-BE3
TEAD motif
ETS motif
pNL-BE3(wt)
pNL-BE3(ΔEm)
pNL-BE3(ΔTm)
F
p = 2.23e-10
p = 1.28e-13
p = 0.69
p = 1.74e-13
Luc activity units
pNL3.2
pNL-BE3(wt)
pNL-BE3(ΔTm)
pNL-BE3(ΔEm)
G
reporter : pNL-BE3
p < 2e-16
p < 2e-16
p < 2e-16
p < 2e-16
p = 8.15e-8
p = 1.21e-12
p < 2e-16
p = 4.43e-13
Luc activity units
venus
TEAD1+YAP
TEAD2+YAP
TEAD3+YAP
TEAD4+YAP
venus
TEAD1+TAZ
TEAD2+TAZ
TEAD3+TAZ
TEAD4+TAZ
FLAG-TEADs
HA-YAP
FLAG-TEADs
HA-TAZ
H
reporter : pNL-BE3
p = 5.05e-8
p = 4.73e-5
p = 0.06
Luc activity units
venus
ELF3
TEAD1+TAZ
TEAD1+TAZ +ELF3
Myc-ELF3
FLAG-TEAD1
HA-TAZ

**Figure S9. TEAD/YAP/TAZ regulation of BE3 activity. A**, BE3 sequence. ETS and TEAD motifs are highlighted in blue and green, respectively. **B**, RNA-seq quantification of *TEAD1*, *TEAD2*, *TEAD3*, *TEAD4*, *YAP1*, and *WWTR1* (encoding TAZ) in ESCs and dox-treated cells (for 2 or 4 days). Data are shown as mean TPM values (two replicates per condition). **C**, Immunostaining confirming the expression and nuclear localization of TEAD1 and YAP/TAZ in dox-treated (2 days) and untreated cells. YAP/TAZ antibody recognizes both YAP and TAZ. Three independent cultures were used for the experiment. Scale bar; 100 µm. **D**, Luciferase assay using a TEAD reporter (pNL-8xFTIIc) and control (pNL3.2). tet-E3 cells were transfected with the indicated reporters and either untreated or treated with dox for 2 days, and NanoLuc activities were measured, normalized to the control, and plotted as mean ± SD (n = 3 independent experiments). Statistical significance was determined by paired t-test. **E**, Schematic of BE3 reporter constructs lacking either the ELF3-binding motif or the three TEAD binding motifs. **F**, Luciferase assay using the indicated reporter constructs. NanoLuc activities were measured 48 h after transfection, normalized to the control, and plotted as mean ± SD (n = 3 independent experiments). Statistical significance was determined by Tukey’s test. **G**, Luciferase assay using the BE3 reporter. Cells were transiently transfected with BE3 NanoLuc reporters plus an internal control reporter and the indicated expression plasmids. NanoLuc activities were measured at 48 h after transfection, normalized to the control, and shown as means ± SD (n = 3 independent experiments). Expressions of tagged TEADs, TAZ, and ELF3 were confirmed by western blotting. Statistical significance was determined by Dunnett’s test (vs Venus-transfected cells). **H**, Luciferase assay using the BE3 reporter was performed as indicated in g. Expressions of tagged TEAD1, TAZ, and ELF3 were confirmed by western blotting. Statistical significance was determined by Tukey’s test. **I**, Representative WB images of co-immunoprecipitation (three independent experiments). Lysates were prepared from day-2 dox-treated cells (HA-ELF3 induced) and subjected to immunoprecipitation with anti-TEAD1 antibody or control IgG. Immunoblots were performed with anti-HA (ELF3), anti-YAP, anti-TAZ, and anti-pan-TEAD antibodies.

Table S1. Target sequences for gRNAs.

| Target | target sequence | PAM |
|---|---|---|
| ELF3, intron 1 | 5'-CTGGGGGCTACTCTTGCCCA-3' | GGG |
| ELF3, intron 7 | 5'-GAGGAAATAAGGCTCCCAGT-3' | GGG |
| BMP4 enhancer (BE3), upstream | 5'-ATACTGGGGGTAGTAGATAA-3' | AGG |
| BMP4 enhancer (BE3), downstream | 5'-GCTCTGGATTCAGACTACAT-3' | GGG |

Table S2. Primer sequences.

Primers for qPCR.

| Application | mRNA | Forward sequence | Reverse Sequence |
|---|---|---|---|
| *qPCR* | *VTCN1* | 5'-GTCGGAGCAGGATGAAATGT-3' | 5'-TAGGTGCCAGCATCTGTGAG-3' |
| | *GABRP* | 5'-ATGGCAGCCAAAGATAGGGG-3' | 5'-TGGCAAAGCTGATCTTCCGT-3' |
| | *SPRR2A* | 5'-CCACCCTGCCAGTCAAAGTA-3' | 5'-TGAAGGTGGAGCTGTGGAAC-3' |
| | *BMP4* | 5'-GGCTGGAATGACTGGATTGT-3' | 5'-TGGTTGAGTTGAGGTGGTCA-3' |
| | *CTGF* | 5'-CAAGGGCCTCTTCTGTGACT-3' | 5'-ACGTGCACTGGTACTTGCAG-3' |
| | *ISL1* | 5'-GTGTGATCCGGGTCTGGTTT-3' | 5'-TTTTGTCATTGGGCTGCTGC-3' |
| | *DLX5* | 5'-GCCAAAGCTTATGCCGACTA-3' | 5'-GCCATTCACCATTCTCACCT-3' |
| | *VGLL1* | 5'-CTGGTCCCATGGCTGTGAAT-3' | 5'-AGGAGAAATGCCACAGCTCC-3' |
| | *GAPDH* | 5'-GAGTCAACGGATTTGGTCGT-3' | 5'-GACAAGCTTCCCGTTCTCAG-3' |
| | Gene locus | Forward sequence | Reverse Sequence |
| genomic PCR | ELF3 | 5'-GGGATGACAGACTCTGACAATCAT-3' | 5'-CTTCTCGTAGGTCATGTTGCTGTTC-3' |
| | BE3 | 5'-CATGTCTCTCCTGAGCATTTGGTTTG -3' | 5'-GAGAAAGAAGTGAGGAAAGTGGTGG-3' |
| | Gene locus | Forward sequence | Reverse Sequence |
| ChIP-qPCR | BE3 | 5'-AAGTCAGTTCCTCCTCTCCCTC-3' | 5'-ACCAGCCAGCATTGTCTCAAC-3' |

Table S3. Antibody list.

| Application | Antibody | Supplier | Cat. Number | Dilution | Note |
|---|---|---|---|---|---|
| IF | TFAP2A | Abcam | Cat# ab52222 | 1/2000 | |
| | CK7 | Dako | Cat# M7018 | 1/200 | |
| | NANOG | R&D Systems | Cat# AF1997 | 1/200 | |
| | OCT3/4 | BD Biosciences | Cat# 611202 | 1/200 | |
| | GATA3 | R&D Systems | Cat# AF2605 | 1/200 | |
| | ISL1 | R&D Systems | Cat# AF1837 | 1/1000 | |
| | VTCN1 | Abcam | Cat# ab252438 | 1/500 | |
| | PRTG | Sigma Aldrich | Cat# HPA032138 | 1/100 | |
| | VGLL1 | Sigma Aldrich | Cat# HPA042403 | 1/2000 | |
| | SDC1 | Proteintech | Cat# 67155-1-Ig | 1/3000 | |
| | HA | Roche | Cat# 11-867-423-001 | 1/1000 | |
| | ELF3 | Sigma Aldrich | Cat# HPA003479 | 1/500 | |
| | pSMAD1/5/8 | Cell Signaling Technology | Cat# 13820 | 1/500 | |
| | Myc | Cell Signaling Technology | Cat# 2278 | 1/1000 | |
| | TEAD1 | BD Biosciences | Cat# 610922 | 1/200 | |
| | YAP | Santa Cruz | Cat# sc-101199 | 1/5000 | also recognize TAZ |
| WB | HA | Cell Signaling Technology | Cat# 2367 | 1/1000 | |
| | ACTIN | Sigma Aldrich | Cat# A5060 | 1/1000 | |
| | ELF3 | Sigma Aldrich | Cat# HPA003479 | 1/1000 | |
| | GABRP | Abcam | Cat# ab26055 | 1/1000 | |
| | IGFBP3 | Protaintech | Cat# 10189-2-AP | 1/1000 | |
| | WNT6 | Abcam | Cat# Ab50030 | 1/2000 | |
| | H3K27ac | Cell Signaling Technology | Cat# 8173 | 1/1000 | |
| | H3K4me3 | Cell Signaling Technology | Cat# 9751 | 1/1000 | |
| | FLAG | Sigma Aldrich | Cat# F3165 | 1/1000 | |
| | Myc | Merck Millipore | Cat# 05-724 | 1/1000 | |
| | TEAD1 | BD Biosciences | Cat# 610922 | - | for IP ; 2 μg |
| | mouse IgG | Proteintech | Cat# 66360-1-Ig | - | for IP ; 2 μg |
| | YAP | Bethyl Laboratories | Cat# A302-309A | 1/1000 | |
| | YAP/TAZ | Cell Signaling Technology | Car# 8418 | 1/1000 | |
| | pan-TEAD | Cell Signaling Technology | Cat# 13295 | 1/1000 | |
| FCM | APA | BD Bioscience | Cat# 564532 | NA | 5 μl |
| | B7-H4-PE | Invitrogen | Cat# 12-5949-42 | NA | 5 μl |
| | B7-H4-APC | Invitrogen | Cat# 17-5949-42 | NA | 5 μl |

| | | | | | |
|---|---|---|---|---|---|
| ChIP | HA | Cell Signaling Technology | Cat# 3724 | NA | 5 µl |
| | H3K27ac | Cell Signaling Technology | Cat# 8173 | NA | 5 µl |
| | H3K4me3 | Cell Signaling Technology | Cat# 9751 | NA | 5 µl |
| | H3K27me3 | Cell Signaling Technology | Cat# 9733 | NA | 5 µl |
| | p300 | Cell Signaling Technology | Cat# 54062 | NA | 5 µl |
| | MED1 | Bechyl Laboratories | Cat# A300-793A | NA | 5 µl |

IF; immunofluorescence, WB; western blot, FCM; flow cytometry)
